# Targeting fungal BET bromodomains as a *pan*-*Candida* antifungal strategy

**DOI:** 10.1101/2023.02.03.527073

**Authors:** Kaiyao Wei, Justin M. Overhulse, Marie Arlotto, Yingsheng Zhou, Nathan J. Dupper, Boris A. Kashemirov, Cécile Garnaud, Gaëlle Bourgine, Muriel Cornet, Gwenaël Rabut, Charles E. McKenna, Carlo Petosa, Jérôme Govin

## Abstract

Small molecules that target one or both bromodomains (BDs) of human BET proteins are intensely studied as potential new therapeutics against cancer, diabetes and other diseases. The BDs of the fungal BET protein Bdf1 are essential for the human fungal pathogen *Candida albicans*, suggesting BET inhibition as a potential antifungal strategy. However, while the inactivation of both Bdf1 BDs is lethal, that of a single BD only modestly affects viability, implying the need to develop antifungal compounds that selectively target both Bdf1 BDs without inhibiting human BDs. Here, we investigate Bdf1 as a potential antifungal target in *Candida glabrata*, an invasive *Candida* species phylogenetically distant from *C. albicans* and of increasing medical concern. We show that Bdf1 BD functionality is essential in *C. glabrata* and identify a phenyltriazine derivative that targets both Bdf1 BDs with selectivity over human BET BDs. We show that human BET BDs can functionally replace Bdf1 BDs in *C. glabrata* and we use the humanized strains to demonstrate on-target antifungal activity of the phenyltriazine compound. Moreover, by exploiting the humanized and parental fungal strains we identified BET inhibitor I-BET726 to have potent antifungal activity against a broad spectrum of *Candida* species, including azole- and echinocandin-resistant clinical *C. albicans* and *C. glabrata* isolates. Crystal structures suggest how to improve the potency and selectivity of these compounds. Taken together, our findings provide compelling support for the development of BET inhibitors as potential pan-*Candida* antifungal therapeutics.

## Introduction

Invasive fungal infections (IFIs) are a major cause of morbidity and mortality, with over 1.5 million deaths estimated annually worldwide^1^. *Candida* species are among the most significant human fungal pathogens, collectively responsible for 800,000 IFIs per year, yielding 40% mortality and a heavy economic burden^2–4^. The incidence of candidemia, the most common form of invasive candidiasis, has seen an alarming increase in recent years^4–6^. In Europe and North America, the *Candida* species that rank first and second in isolation frequency are *C. albicans* and *C. glabrata*, respectively, accounting for ∼70% of all systemic candidiasis^6, 7^. These fungi are ubiquitous human commensals that asymptomatically colonize the majority of the population. However, under certain conditions (diabetes, cancer, immunosuppression, antibiotic therapy, intravenous catheters or long-term hospitalization) they can lead to life-threatening systemic infections. The current repertoire of antifungal drugs severely limits clinical therapies, representing a major global public health problem^8^. The rapidly emerging multidrug-resistant fungal *Candida auris* epitomizes the threat posed by IFIs^9, 10^.

Small-molecule inhibitors targeting chromatin signaling pathways (“epi-drugs”) have recently entered the clinic or are in clinical trials to treat cancer and other diseases^11–13^. Considerable efforts have focused on the Bromodomain and Extra-Terminal (BET) family of proteins, which regulate gene transcription and chromatin organization^14^. In mammals this family comprises four proteins (Brd2, Brd3, Brd4 and Brdt), of which the best studied is Brd4. BET proteins recognize chromatin through their two bromodomains (BDs), BD1 and BD2, which specifically recognize histones acetylated on lysine, including histone peptide sequences bearing two or more closely spaced acetyllysines^15, 16^. Many small-molecule BET inhibitors (BETi) that directly compete with the acetylpeptide ligand have been developed, including several currently in phase II or phase III clinical trials for cancer, diabetes and cardiovascular disease^13, 17^.

Except for certain histone deactylases (HDACs), epigenetic targets have largely remained unexplored in the fungal infection field^18^. One attractive target is the fungal BET protein, Bdf1. Its function as a global transcriptional regulator has best been studied in *S. cerevisiae*^19–21^, where Bdf1 interacts with the transcription factor TFIID^22^ and is a subunit of the SWR1 chromatin remodelling complex^23, 24^. Bdf1 also plays a key role in the salt stress response^25^ and in regulating transcription and chromatin compaction during yeast sporulation^26, 27^. Unlike mammals and *S. cerevisiae*, *Candida* species possess only a single BET gene, *BDF1*^28^. We previously showed that Bdf1 BD functionality is essential in *C. albicans*: mutations inactivating both domains cause a loss of viability *in vitro* and decreased virulence in mice^29^. Interestingly, the BET inhibitor JQ-1 was recently reported to inhibit the growth and virulence of *Aspergillus fumigatus*^30^, suggesting that BET BD functionality is essential in diverse fungal pathogens. In our previous study, we identified small molecules that inhibit individual BDs of *C. albicans* Bdf1 (*Ca*Bdf1) with high selectivity over human BDs. Compounds selective for *Ca*Bdf1 BD1 showed antifungal activity when tested on *C. albicans* strains harbouring inactivating mutations in BD2, and vice versa, establishing *Ca*Bdf1 as a potential new antifungal drug target.

However, identifying compounds with antifungal activity against wildtype clinical reference strains of *C. albicans* through traditional screening and compound optimization approaches has proved challenging. This is largely because of the requirement to selectively target both *Ca*Bdf1 BDs without significantly inhibiting human BDs. The cumbersome nature of genetic manipulations in *C. albicans* has also hampered rapid progress. Furthermore, it is unclear whether a Bdf1-selective dual BD inhibitor would be effective against a broad spectrum of *Candida* species, or whether each species would require the development of its own specific inhibitor, thereby severely limiting usefulness in the clinic.

To address these issues we investigated Bdf1 inhibition in *C. glabrata*. Although in the same genus, *C. glabrata* is phylogenetically, genetically and phenotypically very different from *C. albicans*, more closely resembling *S. cerevisiae* ^31–33^. As the second most frequent cause of invasive candidiasis, *C. glabrata* is of great clinical concern because of its ability to rapidly develop multidrug resistance under antifungal pressure. From a practical perspective, its haploid genome renders genetic manipulations much simpler than in diploid *C. albicans*, while its similarity to *S. cerevisiae* facilitates the transfer of yeast-based molecular technologies.

Here, we show that, as in *C. albicans*, the dual inactivation of both Bdf1 BDs is required for lethality in *C. glabrata*. We identified a phenyltriazine compound that inhibits both Bdf1 BDs selectively over human BET BDs and engineered a *C. glabrata* strain expressing a humanized Bdf1 variant to demonstrate that the compound’s antifungal activity is an on-target effect. The use of this strain additionally allowed us to identify a BET inhibitor, I-BET726, with potent antifungal activity against a wide range of *Candida* species, including drug-resistant clinical isolates. We also report the structural basis for the potency and selectivity of these compounds, providing insights into their chemical optimization. Taken together our findings strongly support the feasibility of developing BET inhibition as a pan-*Candida* antifungal therapeutic strategy.

## Results

### Bdf1 bromodomain functionality is essential in *C. glabrata*

To verify the functionality of BDs in *C. glabrata* Bdf1 (*Cg*Bdf1, **Fig. 1a**), we profiled their ability to bind acetylated histone peptides. We assessed the binding of recombinant *Cg*Bdf1 BD1 and BD2 (*Cg*BD1 and *Cg*BD2) to a commercial array of human histone N-terminal peptides, primarily from histones H3 and H4 (347 of 384 peptides), bearing various post-translational modifications (PTMs). Use of the array is justified because the N-terminal tails of histones H3 and H4 are nearly identical between human and *C. glabrata* (**Fig. S1a**). Like other BET BDs^16, 27, 29^ (**Fig. 1b**), *Cg*Bdf1 BDs recognized several acetylated H3 and H4 peptide sequences, with higher affinity for multi-acetylated peptides and the strongest binding observed towards a tetra-acetylated H4 peptide modified on lysine residues 5, 8, 12 and 16 (abbreviated H4ac4; **Fig. 1c** and **Fig. S1b**). The two BDs exhibit very similar binding profiles: of the 20 acetylated H3 and H4 peptides on the array, 15 showed either significant (14 peptides) or poor (1 peptide) binding by both BDs, whereas only 5 were preferentially bound by either BD1 (4 peptides) or BD2 (1 peptide) (**Fig. 1c**). The observed binding profiles are comparable to those previously reported for *Ca*Bdf1 BD1 and BD2 (*Ca*BD1 and *Ca*BD2)^29^, except that the *Cg*Bdf1 BDs exhibit broader ligand selectivity (**Fig. S1c**). This is especially true for *Cg*BD2, whose binding profile included all acetylpeptides recognized by *Ca*BD2 plus 8 additional peptides. Consequently, the BD1 and BD2 binding profiles are more closely matched for *Cg*Bdf1 than for *Ca*Bdf1 (**Fig. S1c**).

**Figure 1.**
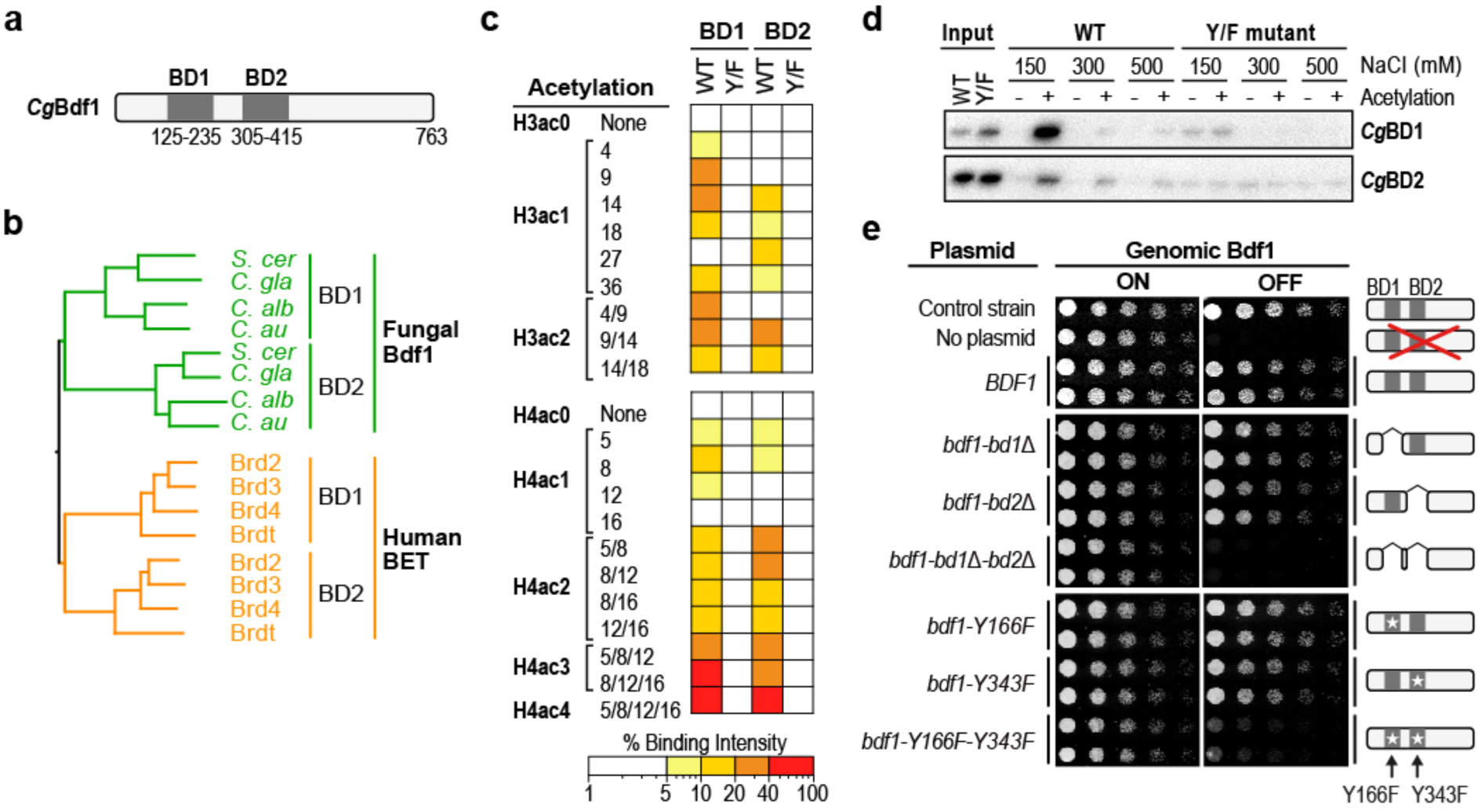
Bdf1 bromodomain functionality is essential in *C. glabrata*. (**a**) Domain boundaries of *Cg*BDF1 BDs. (**b**) Phylogeny of fungal and human BET BDs. (**c**) Summary of binding intensities of WT and mutant forms (Y166F or Y343F) of *Cg*BDF1 BD1 and BD2 to an array of acetylated H3 and H4 peptides. Binding intensities are normalized to a positive control signal, obtained from the interaction of an anti-c-myc antibody with a c-myc control peptide included on the array. (**d**) Pull-down assay. Immobilized H4ac0 and H4ac4 peptides were incubated with GST-tagged *Cg*Bdf1 BDs or with the corresponding Y/F mutants. After washing with buffers containing either 150, 300 or 500 mM NaCl, bound proteins were eluted and visualized by western blotting with an anti-GST antibody. (**e**) Colony formation assays showing the effect of Bdf1 repression or Bdf1 BD inactivation on *C. glabrata* growth.

In the case of *S. cerevisiae* and *C. albicans* Bdf1, BD-mediated ligand binding is disrupted by mutating a conserved tyrosine implicated in acetyllysine recognition to a phenylalanine^20, 21, 27, 29^. Introducing the corresponding Tyr*→*Phe point mutation in *Cg*BD1 and *Cg*BD2 (“Y/F” mutations Y166F and Y343F, respectively) abolished the interaction with acetylated peptides, confirming binding specificity (**Fig. 1c** and **Fig. S1b**). These results were further confirmed in pull-down assays with immobilized H4 peptides. Whereas the wildtype (WT) BDs showed specific H4ac4 peptide binding, those bearing the Y/F mutation bound weakly and failed to discriminate between the acetylated and non-acetylated H4 peptide (**Fig. 1d**). *Cg*BD1 bound the H4ac4 peptide more efficiently than *Cg*BD2 (**Fig. 1d**), reminiscent of the differential binding behaviour of the corresponding mammalian Brd2, Brd4 and Brdt BDs^21, 49^. This differs from the behaviour previously observed for the two *Ca*Bdf1 BDs, which bind the H4ac4 peptide with comparable strength^29^. Hence, the broader ligand selectivity observed for *Cg*BD2 compared to *Ca*BD2 appears to be at the expense of H4ac4 binding affinity.

To gain additional insights, we solved the crystal structures of *Cg*BD1 and *Cg*BD2 (**Suppl. Table 1**). These adopt the expected BD fold defined by helices Z, A, B and C and closely resemble the structures of human BET and *Ca*Bdf1 BDs (**Fig. S2a and b**; **Suppl. Table 2**). The *Cg*BD1 and *Cg*BD2 ligand binding pockets, defined by the ZA and BC loops, are structurally well conserved, including all water molecules implicated in ligand recognition ^17, 34^ (**Fig. S2c and d**). Structures reported for mammalian BET BDs bound to mono- and di-acetylated histone peptides reveal a binding mode in which the unique or primary acetyllysine (Kac) binds deep within the BD’s hydrophobic pocket while flanking residues bind along a shallow groove in a common N-to-C peptide orientation. The ensemble of co-crystallized peptides contact a total of 20 BD residues located in helices B and C and in the ZA and BC loops, including 13 residues that interact with one or both Kac residues (**Fig. S3** and **S4**). The corresponding 13 positions are invariant or highly conserved between *Cg*BD1 and *Cg*BD2 (**Fig. S3a**), as well as between orthologous *Cg*Bdf1 and *Ca*Bdf1 BDs (**Fig. S3a**), explaining the similar ligand selectivities of these BDs. Structural alignments suggest that a small number of divergent residues at these positions could feasibly account for the binding differences observed between *Cg*BD1, *Cg*BD2 and *Ca*BD1 (**Fig. S3b-e** and **Fig. S4b,d**) and for the comparatively narrower range of acetylated peptides recognized by *Ca*BD2 (**Fig. S4c,e**).

To verify whether Bdf1 BD functionality is essential in *C. glabrata*, we replaced the endogenous *BDF1* promoter by that from the *MET3* gene (*pMET*), which is inhibited by adding an excess of methionine and cysteine to the growth medium^50^ (**Fig. S5a**). Immunoblotting confirmed that *pMET-BDF1* was effectively repressed by methionine and cysteine, with most Bdf1 protein disappearing within 8 hours of repression (**Fig. S5b,c**). Colony formation assays showed that *BDF1* repression was lethal and that the phenotype was rescued by ectopic expression from an autonomous plasmid, confirming that Bdf1 is essential for *C. glabrata* growth *in vitro* (**Fig. 1e** and **Fig. S5e**). We next generated Bdf1 mutants compromised for ligand-binding activity by deleting one or both BDs or introducing one or both Y/F point mutations. The introduction of these mutations did not significantly affect Bdf1 expression levels (**Fig. S5d**). Whereas disrupting only a single BD did not markedly affect growth, simultaneously disrupting both was lethal (**Fig. 1e, Fig. S5e**), demonstrating that the presence of at least one functional BD within Bdf1 is required for the viability of *C. glabrata in vitro*.

### BET inhibitors can discriminate between human and *C. glabrata* BET BDs with high selectivity

The similar acetylpeptide binding profiles of human BET and *Cg*Bdf1 BDs underscore the conserved nature of their ligand binding pockets. However, for BET inhibition to be a feasible antifungal strategy, these pockets must sufficiently differ to allow small-molecule inhibitors to selectively target the fungal BDs. Indeed, our *Cg*Bdf1 BD crystal structures reveal differences with those of human BET BDs, suggesting the potential for selective inhibitor development. To gain further insights, we used commercially available BET inhibitors as probes to assess how well small molecules could discriminate between human BET and *Cg*Bdf1 BDs. Specifically, we evaluated the ability of BETi compounds JQ1, I-BET151, bromosporine and PFI-1 to inhibit BDs from binding an H4ac4 peptide using a homogeneous time-resolved fluorescence (HTRF) assay^29^ adapted for *Cg*Bdf1 BDs (**Fig. 2a**). As expected, the four compounds potently inhibited human Brd4 BD1 and BD2, yielding median inhibitory concentrations (IC_50_ values) of ∼10-100 nM (*p*IC_50_ values of 7-8), resembling previously reported values ^35–37^ (**Fig. 2b**). In contrast, the *Cg*Bdf1 BDs were more weakly inhibited, with IC_50_ values between 0.5 and >100 μM (*p*IC_50_ values of <4-6.3), revealing a reduction in sensitivity of approximately 1 to 3 log (*Δp*IC_50_ = 1 to 3; **Fig. 2c**). The BETi sensitivity of both *Cg*BDs followed the order I-BET151 > JQ1 > Bromosporine > PFI-1 in decreasing order of potency, with *Cg*BD2 consistently more sensitive (by at least 4-fold) than *Cg*BD1 to all compounds except PFI-1, which weakly inhibited *Cg*BD1 but not *Cg*BD2. Not surprisingly, none of these BETi compounds significantly inhibited the growth of *C. glabrata in vitro* (**Fig. 2d**).

**Figure 2.**
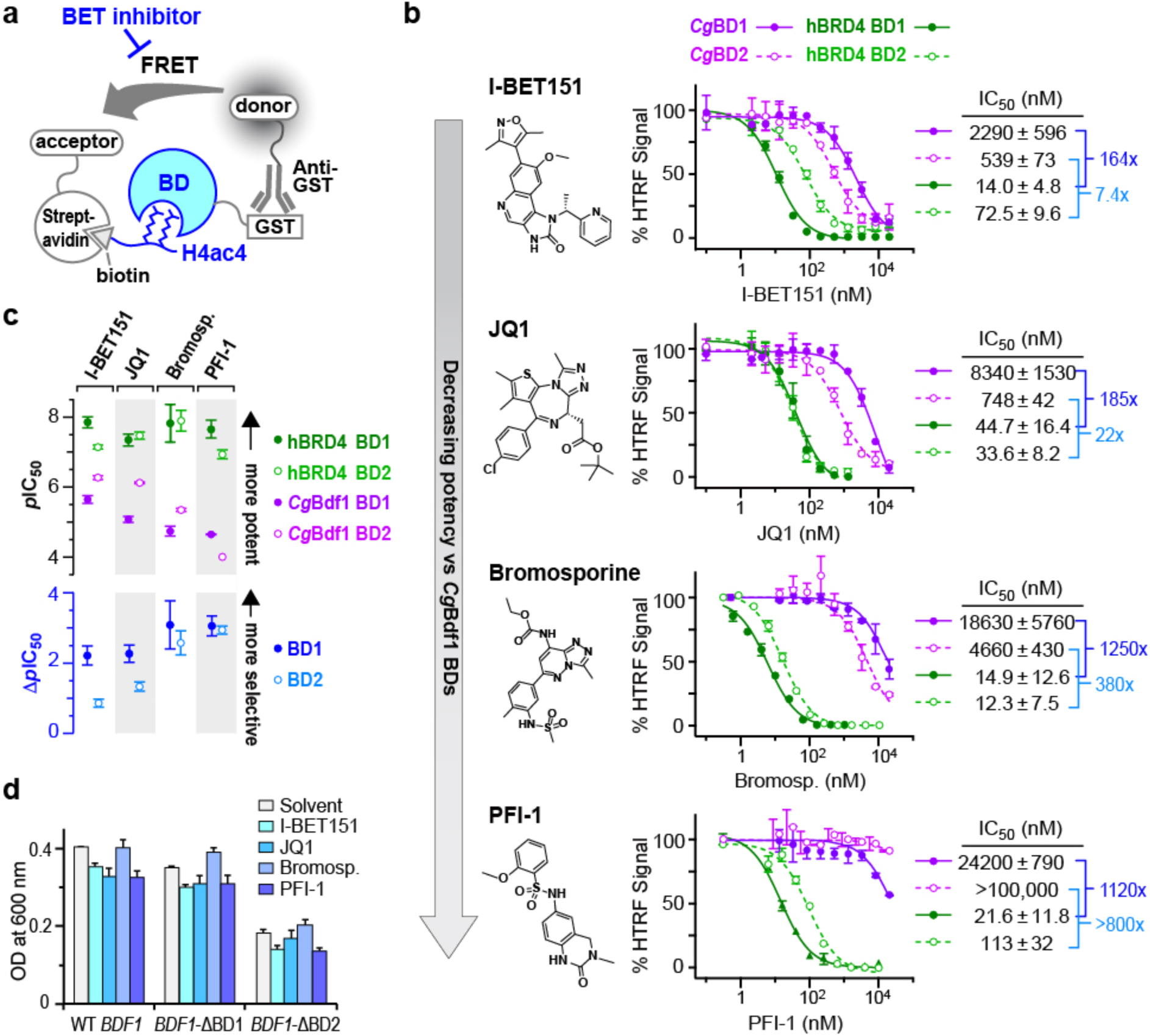
BET inhibitors can discriminate between human BET and *Cg*Bdf1 BDs with high selectivity. (**a**) Overview of the HTRF assay. A GST-tagged BET BD is bound to an anti-GST antibody coupled to the donor fluorophor (Lumi4-Terbium cryptate). A biotinylated H4ac4 peptide is bound to streptavidin coupled to the acceptor fluorophore (D2 dye). The binding of GST-BD to H4ac4 brings the donor and acceptor into close proximity, allowing for FRET. Following donor excitation, fluorescence emitted by the acceptor is detected following a delay between the excitation pulse and the time-gated measurement window. The presence of a BET inhibitor disrupts the binding interaction, resulting in reduced HTRF signal. (**b**) HTRF assays performed on *Cg*Bdf1 BD1 and BD2 (closed purple and open magenta circles, respectively) and human Brd4 BD1 and BD2 (closed dark green and open light green circles, respectively) in the presence of the indicated compound. IC_50_ values are listed next to each graph. Data represent the mean and s.d. values from three independent experiments. (**c**) Potency and selectivity of BETi compounds. *Top.* Summary of *p*IC_50_ values showing the potency of compounds against Brd4 and *Cg*Bdf1 BDs. Symbol definitions are as in (b). *Bottom.* Summary of *Δp*IC_50_ values showing the selectivity of inhibitors towards the human BDs. Closed blue (BD1) and open cyan (BD2) circles represent the difference in *p*IC_50_ value between corresponding Brd4 and *Cg*Bdf1 BDs. (**d**) The four BET inhibitors do not affect *C. glabrata* growth, even when Bdf1 BD1 or BD2 is deleted. Expression of endogenous *BDF1* was suppressed by a 23h incubation with methionine and cysteine and hence only the WT or mutant Bdf1 protein encoded by the plasmid was expressed. Strains were then incubated with BETi compounds at 10 μM for 24h and viability was assessed. Data represent the mean and s.d. values from three independent experiments.

To better understand these results we sought crystal structures of BETi-bound *Cg*BDs but only succeeded in crystallizing *Cg*BD2 in complex with I-BET151, the *Cg*BD/BETi combination with the highest potency of inhibition (**Fig. 3a**). The structure of this complex closely resembles that of unbound *Cg*BD2, with one notable difference. In the unbound structure the side chain *χ*1 angle of Tyr327 in the “YPF-shelf” is stabilized in a *gauche^-^* conformation by a sulfur-*π* interaction with residue Met395. I-BET151 induces a switch to the *trans* conformation, such that the Tyr327 side chain flips away from the binding pocket, relieving the steric clash that would otherwise occur with the ligand (**Fig. 3b**).

**Figure 3.**
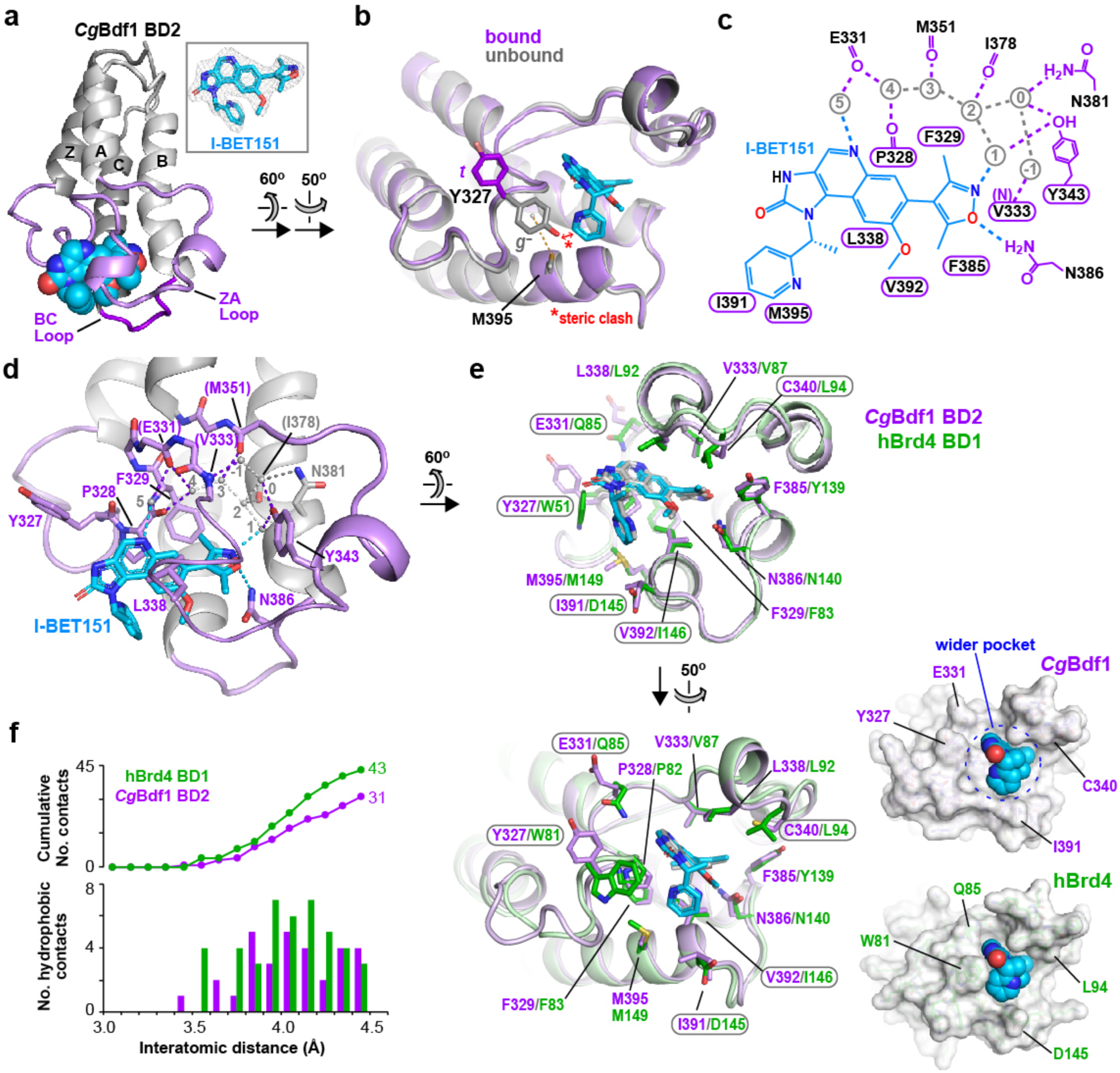
A wider binding pocket explains the reduced potency of I-BET151 towards *Cg*BD2. (**a**) Crystal structure at 1.95 Å resolution of *Cg*BD2 bound to I-BET151. *Inset.* Simulated annealing omit *F*_o_- *F*_c_ density for I-BET151 contoured at 3.5 *σ*. (**b**) Structural alignment of *Cg*BD2 in the unbound state (grey) and bound to I-BET151 (violet). The two aligned structures have an rmsd of 0.418 Å for 108 C*α* atoms. The dashed line shows the sulfur-*π* interaction involving residues Tyr327 and Met395. The ligand-induced switch in Tyr327 side chain conformation involves a change of *χ*1 angle from a *gauche^-^* (*g*^-^) to *trans* (*t*) configuration. This switch is not due to crystal packing effects because the *gauche^-^* and *trans* Tyr327 conformations are consistently observed in all seven and all four *Cg*BD2 molecules in the asymmetric units of the respective crystal forms. (**c**) Schematic summary of ligand interactions. Hydrogen bonds are shown as dashed lines. Residues mediating van der Waals contacts with I-BET151 are indicated by labels within a cartouche. Water molecules are indicated in grey and are numbered as in ref. ^53^. (**d**) Conserved water structure and hydrogen bonding interactions in the binding site. Residues interacting with I-BET151 through direct and water-mediated hydrogen bonds (dashed lines) are shown in stick representation. Residues interacting through backbone atoms are labelled in parentheses. (**e**) Alignment of the I-BET151-bound structures of *Cg*BD2 (violet) and Brd4 BD1 (PDB 3ZYU; green). The ligands bound to *Cg*BD2 and to Brd4 BD1 are shown in cyan and grey, respectively. Labels within a cartouche indicate *Cg*BD2 residues that contact I-BET151 less intimately than the corresponding Brd4 BD1 residue. *Bottom left.* Surface representations showing the wider binding pocket of *Cg*BD2 (top) compared to Brd4 BD1 (bottom). (**f**) Number of hydrophobic contacts between BD atoms and I-BET151 plotted as a function of interatomic distance, shown as a frequency histogram (bottom) and cumulative frequency distribution (top). Hydrophobic contacts were identified using the web server Arpeggio ^54^.

I-BET151 adopts the same pose as previously observed with the BD1 domains of human Brd2 and Brd4 ^36, 38^ and preserves the corresponding hydrogen bonding interactions, including a direct H bond with Asn386 and two water-mediated H bonds with the Tyr343 hydroxyl and the Glu331 main chain carbonyl (**Fig 3c,d**). However, it interacts less intimately with the *Cg*BD2 binding pocket than with that of human BDs. This primarily reflects the lack of contact between the *trans* conformation of Tyr327 and I-BET151, whereas the corresponding WPF-shelf Trp residue in human BET BDs (Trp51 in Brd4) packs against the pyridine and imidazoquinoline rings (**Fig. 3e**). The fact that certain *Cg*BD2 residues have shorter side chains (Cys340, Val392) or an altered conformation (Glu331) compared to their Brd4 counterparts also contributes to the more open appearance of the ligand binding pocket (**Fig. 3e**). Consequently, compared to Brd4-BD1, *Cg*BD2 residues mediate fewer hydrophobic contacts with I-BET151, especially at interatomic distances below 4 Å (**Fig. 3f**). This difference is reflected by the solvent-accessible surface area (SASA) of I-BET151 buried by each BD: of the ligand’s total SASA (3646 Å^2^), 55.4% is buried in the Brd4 binding pocket compared to only 47.2% in that of *Cg*BD2. Similarly, a structural alignment of I-BET151-bound *Cg*BD2 with that of unbound *Cg*BD1 predicts that the *Cg*BD1 binding pocket would engage I-BET151 less intimately (burying only 43.4% of the ligand’s total SASA) than Brd4 BD1. This difference is primarily due to *Cg*BD1 residues Arg150 and Gln154 pointing outward away from the ligand, unlike the corresponding inward-pointing Trp81 and Gln85 residues of Brd4 (**Fig. S6a**). The looser fit of I-BET151 within the *Cg*BD1 and *Cg*BD2 binding pockets likely explains the decreased potency observed for this compound relative to the human BDs. The more open BD binding pockets of *Cg*Bdf1 is reminiscent of a similar observation previously reported for *C. albicans* Bdf1 ^29^.

We next modelled the structures of *Cg*BD1 and *Cg*BD2 bound to JQ1, bromosporine and PFI-1 by aligning the corresponding ligand-bound structures of Brd4 BD1 and Brd2 BD2 with our available *Cg*BD1 and *Cg*BD2 crystal structures (**Fig. S6**). None of these models yielded a major steric clash between the *Cg*BD and superimposed BETi ligand. Instead, as observed for I-BET151, the three BETi compounds exhibited a relatively loose fit into the *Cg*BD binding pockets, on average burying ∼200 Å^2^ less SASA and making fewer contacts than in the corresponding human structures (**Fig. S6b-d**). In the JQ1- and PFI-1-bound models, the poorer fit primarily arises from the same structural differences between the human and fungal binding pockets as described for I-BET151. An additional difference contributes to the poor fit of BSP: in the human BET BDs the amino group of a lysine (Lys91 in Brd4 BD1 and Lys374 Brd2 BD2) is within H-bonding distance of the BSP sulfonamide group, whereas the corresponding *Cg*BD1 (Ala160) and *Cg*BD2 (Glu331) side chains are either too short or lack the required proton to mediate such an interaction (**Fig. S6c**). Taken together, these results demonstrate that small molecules can discriminate the human BET and *Cg*Bdf1 BDs with high selectivity.

### Chemical screening identifies *Cg*BD1-selective compounds and non-selective dual *Cg*BD inhibitors

We next adapted our HTRF assay to perform a high-throughput (HT) screen of small molecule inhibitors selective for *Cg*Bdf1 BDs over human BET BDs. Because the HT assay parameters (signal-to-background ratio and Z’-factor) were more favorable for *Cg*BD1 than *Cg*BD2, we performed the primary screen against *Cg*BD1. Screening a library of ∼100,000 chemically diverse compounds resulted in ∼250 confirmed hits, of which 63 yielded median inhibitory concentration (IC_50_) values in the low micromolar range and were commercially available. We repurchased these compounds, including 35 distinct scaffolds (**Fig. S7**), and rescreened them in our initial (low-throughput) HTRF assays (**Suppl. Table 3**). The 12 most potent hits (10 scaffolds) were then further tested in multi-replicate HTRF assays with *Cg*BD1 and *Cg*BD2 and the corresponding human Brd4 BDs (**Suppl. Table 4**). This revealed two groups of compounds: those that selectively inhibited *Cg*BD1 (8 compounds) and those that additionally inhibited *Cg*BD2 as well as both Brd4 BDs (4 compounds) (**Fig. 4a**).

**Figure 4.**
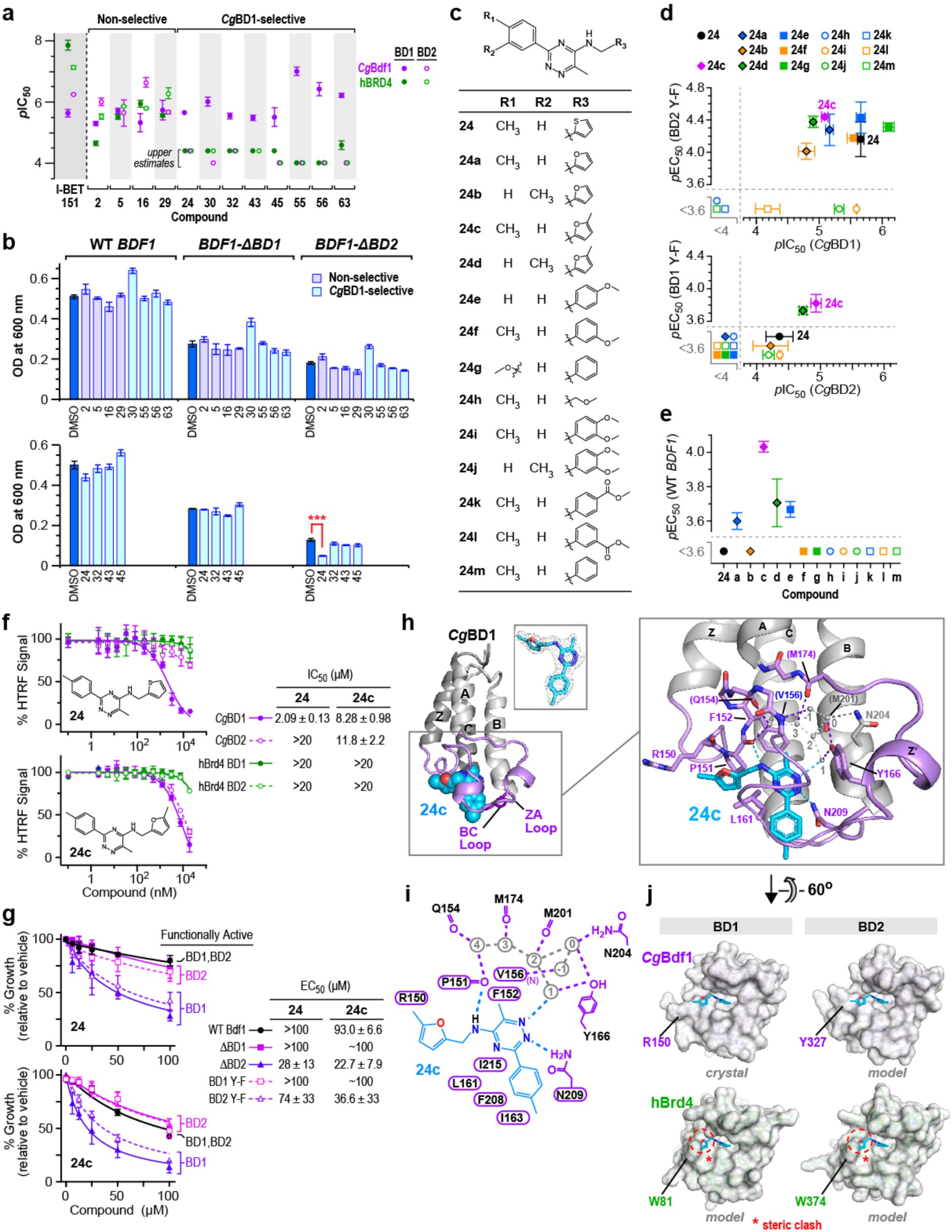
Identification of *Cg*Bdf1-selective compound 24c with dual BD inhibitory activity. (**a**) Summary of HTRF assay results for 12 selected HTS hits. The *p*IC_50_ values are shown for *Cg*BD1 and *Cg*BD2 (closed purple and open magenta circles, respectively) and human Brd4 BD1 and BD2 (closed dark green and open light green circles, respectively). See also **Suppl. Table 4**. (**b**) Summary of growth inhibition assays against *C. glabrata* strains expressing WT and mutant Bdf1. (**c**) Analogs of **24** used in this study. (**d,e**) Inhibitory activity of analogs of **24** towards single Bdf1 BDs. Inhibition observed in HTRF assays with *Cg*BD1 and *Cg*BD2 is compared with growth inhibition observed with *C. glabrata* strains expressing Bdf1 mutated in the other BD. *Top*. *p*IC_50_ values determined in HTRF assays with *Cg*BD1 plotted against *p*EC50 values determined in growth inhibition assays on a strain expressing a BD2-inactivated Bdf1 mutant. *Bottom*. *p*IC_50_ values determined for *Cg*BD2 plotted against *p*EC50 values determined from a strain expressing a BD1-inactivated Bdf1 mutant. Data represent the means ± SD for *n* ≥ 2 independent experiments. (**f**) HTRF inhibition assays performed on **24** (top) and **24c** (bottom) against the BD1 and BD2 domains from *Cg*Bdf1 and human Brd4. (**g**) Growth inhibition assays performed on **24** (top) and **24c** (bottom) with *C. glabrata* strains expressing WT and mutant Bdf1. The BDs functionally active in each strain are indicated. (**h**) Crystal structure of *Cg*BD1 bound to **24c**. *Small inset.* Simulated annealing omit *F*_o_-*F*_c_ density for **24c** contoured at 3 *σ*. *Large inset.* Details of the ligand binding site. Hydrogen bonds are shown as dashed lines. Water molecules are numbered as in ref. ^53^. (**i**) Schematic summary of ligand interactions. Residues mediating van der Waals contacts with **24c** are indicated by labels within a cartouche. Hydrogen bonds and water molecules are shown as dashed lines and grey circles, respectively. (**j**) Surface representations of the *Cg*BD1/**24c** crystal structure and structural alignment models of *Cg*BD2 and of Brd4 BD1 and BD2 showing observed or predicted interactions with the binding pocket. The steric clashes predicted between **24c** and Brd4 residues Trp81 and Trp374 are indicated by red circles and asterisks.

To better understand the stereochemical requirements for selectivity and dual BD activity, we determined BD co-crystal structures for a representative member of each group, non-selective **29** and *Cg*BD1-selective **63** (**Fig. S8a**,**b**). The structure of **29** bound to *Cg*BD2 reveals that the compound is primarily recognized through its acetylpyridoindole moiety. The acetyl group of this moiety mimics that of the histone acetyllysine ligand by forming direct and water-mediated H bonds with Asn386 and Tyr343 in the BC and ZA loops, respectively, while the compound’s morpholinylphenyl moiety packs against Tyr327 (**Fig. S8c**,**d**). *Cg*BD2 adopts the same structure as in the unbound form: in contrast to I-BET151, it binds **29** without flipping the Tyr327 side chain. Structural alignments with the unbound forms of *Cg*BD1 and of human Brd4 BD1 and BD2 show that these BDs all have enough space in their binding pockets to accommodate **29** (**Fig. S8e**), explaining the poor selectivity observed for this compound. The replacement of Tyr327 by the bulkier Trp81 and Trp370 residues in the human BDs would cause a steric clash with the **29** phenyl group, but this could easily be relieved by a small rotation of the compound without disrupting the intimate contacts at the opposite end of the molecule.

*Cg*BD1 recognizes **63** by direct and water-mediated H bond interactions involving the pyrazole 2-nitrogen and residues Asn209 and Tyr166, respectively, as well as a direct H bond between the compound’s carboxyamide nitrogen and the Pro151 carbonyl (**Fig. S8f,g**). The oxadiazole and two thiene moieties pack against residue Met218 within a shallow groove defined by residues Lys147 and RPF-shelf residue Arg150 on one side and Val214 and Ile215 on the other. A structural alignment with *Cg*BD2 shows that these moieties would extensively overlap with Tyr327 in its preferred *gauche^-^* conformation and, unlike I-BET151, topologically prevent its flipping to the *trans* form to accommodate the ligand (**Fig. S8h**), explaining the inability of **63** to inhibit *Cg*BD2. Similarly, a severe steric clash predicted between these moieties and WPF-shelf residues Trp81 and Trp374 explains the weak activity of **63** against the two Brd4 BDs (**Fig. S8h**). These observations suggest that the WPF-shelf Trp residues conserved across human BET BDs may conveniently be exploited to increase the selectivity of compounds towards *Cg*BDs.

### Identification of a *Cg*Bdf1-selective compound with dual BD inhibitory activity

We next tested the 12 inhibitors for their ability to inhibit *C. glabrata* growth in liquid media. Compounds were tested for inhibitory activity against strains expressing WT Bdf1 or a mutant form deleted for either BD1 or BD2. Because the functionality of either BD is sufficient for viability (**Fig. 1e**), dual *Cg*BD inhibitors that successfully target Bdf1 within the fungal cell should inhibit the growth of all three strains, whereas *Cg*BD1-selective compounds should only inhibit the strain expressing Bdf1-*Δ*BD2. Almost none of the compounds showed any significant antifungal activity, suggesting that cellular permeability barriers, extrusion by efflux pumps or other mechanisms prevented them from efficiently inhibiting intracellular Bdf1 (**Fig. 4b**). The one notable exception was **24**. This *Cg*BD1-selective compound inhibited the growth of the strain expressing Bdf1-*Δ*BD2 but not that expressing Bdf1-*Δ*BD1 or the WT protein, suggesting an on-target effect.

We next obtained analogs of **24** (either commercially available or synthesized in-house) that varied at three substituent positions on the phenyltriazine scaffold (**Fig. 4c**) and tested these in HTRF and fungal growth inhibition assays (**Fig. 4d-f**). Shifting the methyl group from the *para* to the *meta* phenyl ring position caused an approximately 2-fold drop in potency against *Cg*BD1 in HTRF assays and a loss of growth inhibition of the Bdf1-*Δ*BD1 expressing strain (**Fig. 2d**, upper panel, compare compounds **24a**, **c** and **i** with **24b**, **d** and **j**, respectively). Although replacing the thiophenyl by a furyl (**24a**) or methylfuryl (**24c**) group caused a 3- or 4-fold drop in *Cg*BD1 inhibition, respectively, it resulted in slightly enhanced growth inhibition of the *BDF1-ΔBD1* mutant, possibly because of improved solubility. In contrast, replacement by a phenyl (**24m**) or methylcarboxyphenyl (**24k** and **l**) group severely compromised all inhibitory activity, whereas replacement by a methoxyphenyl (**24e** and **f**) or dimethoxyphenyl (**24i** and **j**) did not affect the IC_50_ for *Cg*BD1, although the latter replacement abolished growth inhibition. None of the analogs significantly enhanced the potency against human Brd4 BD1 or BD2, indicating that the chemical modifications introduced did not compromise selectivity. Surprisingly, unlike **24** and the other analogs, methylfuryl-containing compounds **24c** and **d** significantly inhibited *Cg*BD2 in the HTRF assay and inhibited the growth of *C. glabrata* strains expressing WT Bdf1 and the Bdf1-*Δ*BD1 mutant (**Fig. 4d**, **e**). Of the two compounds, **24c** exhibited the more potent BD inhibition (IC_50_ values of 2 and 8 μM against *Cg*BD1 and *Cg*BD2, respectively) and antifungal activity against the WT strain (EC50 value of 90 μM) (**Fig. 4f**,**g**).

We next determined the crystal structure of *Cg*BD1 bound to **24c**. This revealed that the methyltriazine moiety occupies the acetyllysine binding site, mediating direct and water-mediated H bonds with BC loop residue Asn209 and ZA loop residue Tyr166, respectively (**Fig. 4h, i**). The *p-* methylphenyl moiety is recognized by hydrophobic residues Leu161, Ile163, Phe208 and Ile215, including a close contact between the *p-*methyl group of **24c** and the Ile163 methyl group. Shifting the compound’s methyl group to a *meta* position would abolish this contact and (in one of the two *meta* positions) yield a steric clash with Phe208, explaining the loss of potency for compounds with this configuration. The methlyfuryl ring rests against Pro151 in the RPF shelf, while the methyl group makes a van der Waals contact with Arg150. Considerable space surrounds this moiety in the binding pocket, explaining why its replacement by certain bulkier substituents is well tolerated.

An alignment with the IBET151-bound structure of *Cg*BD2 suggests that the *Cg*BD2 binding pocket can accommodate **24c** and mediate nearly identical interactions as those mediated by *Cg*BD1 (**Fig. 4j**). Interestingly, superimposing the structures of **24** and **24a** onto that of **24c** reveals that the thiophenyl, furyl and methylfuryl groups of these compounds would all clash with the *gauche-* conformation preferred by Tyr327 in the unliganded state, but that only the methylfuryl group would form a favourable contact with the Tyr327 *trans* conformation, possibly explaining why compounds **24c** and **24d**, but not the other analogs, inhibit *Cg*BD2. The selectivity observed for **24c** for *Cg*BDs over human Brd4 BDs can be attributed to differences in the (R/Y/W)PF shelf in a similar way as described above for **63**. Namely, whereas the side chains of *Cg*BD residues Arg150 and Tyr327 point away from the binding pocket, the corresponding Trp81 and Typ374 side chains of Brd4 point into the pocket and would clash sterically with the compound’s methylfuryl group (**Fig. 4j**). The identification of **24c** demonstrates the feasibility of developing antifungal compounds that target both *Cg*BDs with selectivity over the human BET BDs.

### Humanized *C. glabrata* strains reveal on-target activity of 24c and potent inhibition by I-BET726

We next sought to verify that **24c** inhibited *C. glabrata* growth by engaging Bdf1 BDs in the cell rather than through an off-target effect. To this end we generated *C. glabrata* strains expressing “humanized” variants of Bdf1 (either untagged or C-terminally FLAG-tagged to facilitate immunodetection), in which one or both Bdf1 BDs were replaced by the corresponding human Brd4 BDs (**Fig. 5a**). Growth assays showed that the double BD replacement did not compromise viability and that the humanized strains grew with similar kinetics to the corresponding WT strain (**Fig. 5b**), indicating that the human BDs could functionally replace their fungal counterparts. An immunoblot of cell extracts revealed that humanized Bdf1 was more highly expressed than the WT protein, presumably to compensate for a loss in functional efficacy (**Fig. 5c**).

**Figure 5.**
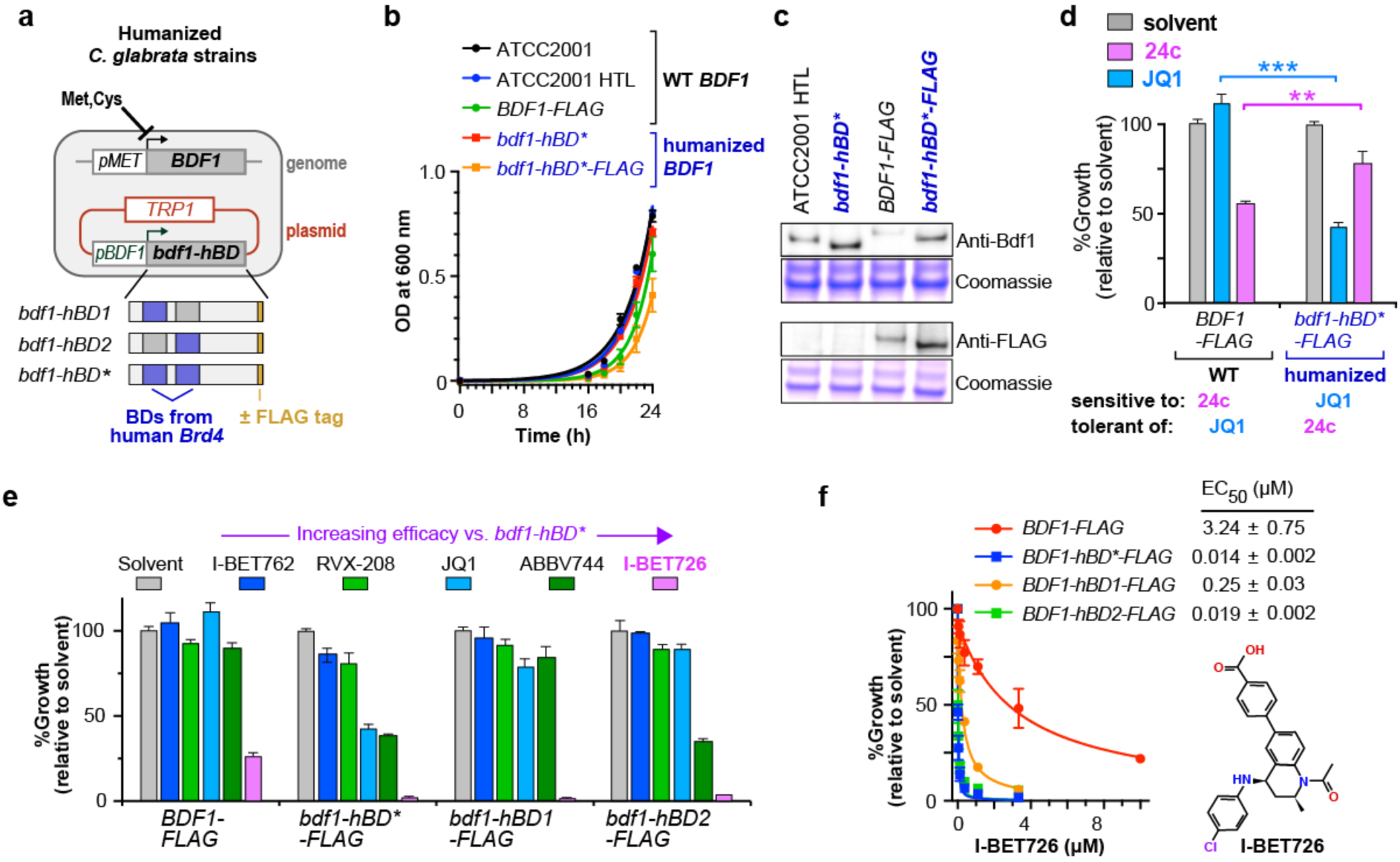
Humanized *C. glabrata* strains reveal on-target activity of 24c and potent inhibition by I-BET726. (a) *Cg*Bdf1 containing BD1 and BD2 from human Brd4 is expressed on a plasmid while expression of the endogenous Bdf1 is repressed by including methionine and cysteine in the growth medium. (b) Growth curves of *C. glabrata* strains expressing WT or humanized Bdf1. (**c**) Western blots showing the expression of WT and humanized Bdf1. Coomassie staining shows equivalent protein loading. (**d**) Growth assays showing the differential sensitivity of *C. glabrata* strains expressing WT and humanized Bdf1 to **24c** and **JQ1.** Compounds were tested at 50 μM concentration. (**e**) Ability of five BET inhibitors tested at 50 μM to inhibit the growth of *C. glabrata* strains expressing WT and humanized Bdf1. (**f**) **I-BET726** inhibits the growth of WT and humanized *C. glabrata* strains.

We then examined the effect of the human BET inhibitor JQ1 and **24c** on the growth of the WT and humanized strains. Notably, JQ1 more potently inhibited the humanized strain than the WT strain, whereas the converse was true for **24c**, mirroring the efficacy of these inhibitors towards human Brd4 and *Cg*Bdf1 BDs *in vitro* (**Fig. 5d**). The results show that JQ1 and **24c** can interact with their intracellular BD targets long enough to compromise viability before being extruded or metabolically inactivated by the fungal cell. These findings strongly confirm that the antifungal activity of **24c** is an on-target BET BD inhibitory effect.

We then took advantage of the humanized Bdf1 strain to test commercially available BETi compounds, including pan-BET inhibitors JQ1, I-BET726 and I-BET762 and BD2-selective molecules RVX-208 and ABBV-744. The different BET inhibitors showed different efficacy against the humanized strains, presumably reflecting differences in fungal cell permeability, efflux rate, and metabolism (**Fig. 5e**). Strikingly, one compound, I-BET726, showed exceptional activity: it markedly diminished the growth of the WT strain and totally abolished growth when either Bdf1 BD was replaced by the corresponding human Brd4 BD (**Fig. 5e**). Dose-response assays revealed EC50 values for I-BET726 of 3 µM and 14 nM against strains expressing WT or humanized Bdf1, respectively (**Fig. 5f**). Thus, I-BET726 appears especially efficient at penetrating the yeast cell and associating with its BD target.

### NanoBiT assays confirm the on-target activity of 24c and I-BET726

We next sought to confirm on-target inhibition by **24c** and I-BET726 by developing an orthogonal assay that probed the histone binding activity of Bdf1 in yeast cells, exploiting *S. cerevisiae* as a convenient surrogate for *C. glabrata* given the many similarities between these species^31–33^. We chose the nanoBiT bioluminescence assay since it has successfully been used to quantify protein-protein interactions in yeast^39^, with significant improvement over other protein complementation assays^40^. This method exploits a luciferase (nanoLuc) that is split into a large (LgBiT, 18 kDa) and a small (SmBiT, 11-residue peptide) fragment that are separately fused to the two interacting proteins of interest (**Fig. 6a**). Upon interaction, the fused moieties can reconstitute an active luciferase that emits light upon addition of the substrate fumirazine. This assay requires the proper physical association of the two luciferase fragments, which is sensitive to the interaction geometry of the fused proteins. For this reason, we screened different histones for the optimal pairing with Bdf1. We took advantage of the SWAT-C collection of *S. cerevisiae* strains^41, 42^ to screen Bdf1 in combination with 10 yeast histone genes (all those in yeast except for the centromeric histone Cse4) as well as other expected Bdf1 interactants, including Bdf2, the H2A.Z chaperone and the 13 subunits of the Swr complex^27^ (not shown). A total of 50 strains expressing nanoBiT tagged proteins were generated with both types of LgBiT/SmBiT pairs. Several strains generated a significant and reproducible nanoBiT signal, including strains expressing nanoBiT-tagged Yaf9, previously shown to mediate the interaction of Bdf1 with the Swr complex^27, 43^ (not shown).

**Figure 6.**
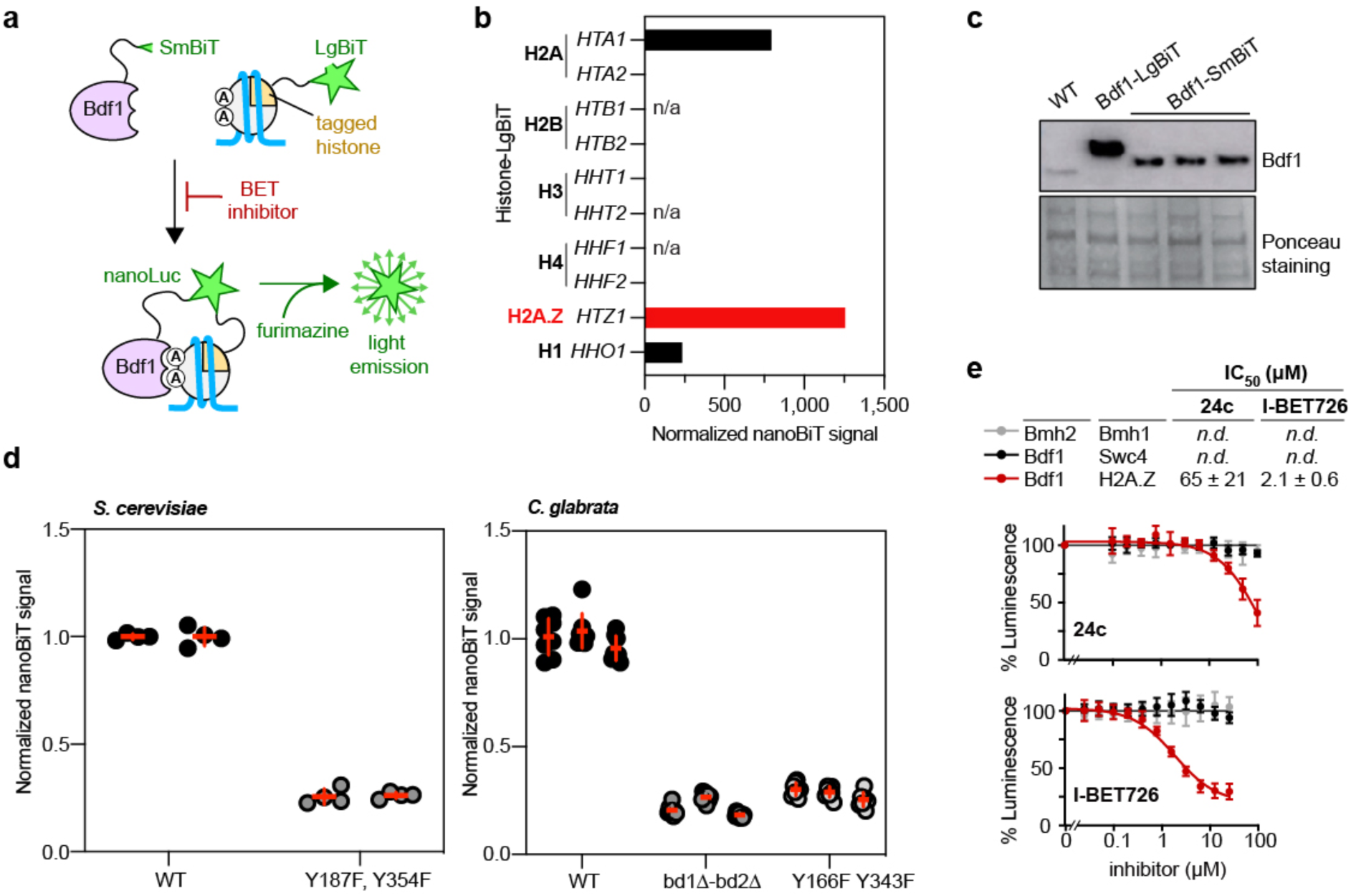
NanoBiT assays confirm the on-target activity of 24c and I-BET726. (**a**) Principle of the nanoBiT assay. A nanoLuciferase is divided into two parts, SmBiT and LgBiT, which are fused to Bdf1 and a histone gene, respectively. Upon recruitment of Bdf1 to chromatin, nanoLuc is reassembled and emits light in the presence of its substrate, furimazine. (**b**) Selection of the best Bdf1/histone gene pair. Most of the histone genes were tagged with LgBiT. H2A.Z (*HTZ1* gene) provided the best signal, probably because it is enriched at the same loci as Bdf1, on the +1 nucleosome (Bdf1 interacts significantly with SWR1C, the complex that loads H2A.Z onto chromatin at the +1 nucleosome). Results shown are those of 2-3 biological replicates (independent yeast isolates), each performed as 3-5 technical replicates. (**c**) Expression levels of Bdf1 when tagged with SmBiT or LgBiT. Bdf1-SmBiT is expressed at levels closer to those of the WT protein. (**d**) The nanoBiT signal is dependent on the integrity of Bdf1 BDs. Bdf1-SmBiT was overexpressed from a plasmid, while H2A.Z was endogenously tagged with LgBiT. When expressing *Sc*Bdf1, the nanoBiT signal is greatly diminished upon mutation of Bdf1 BDs (left panel). Similar results were obtained when *Cg*Bdf1 was expressed in *S. cerevisiae* (right panel). Results shown are those of 2-3 biological replicates (independent yeast isolates), each performed as 3-5 technical replicates. (**e**) *In cellulo* validation of the on-target effect of compounds **24c** and iBET726. Both compounds inhibit nanoBiT signaling in a dose-dependent manner, which is specific for the Bdf1-SmBiT/H2A.Z-LgBiT pair. Bdf1/Swc4 and Bmh1/Bmh2 are known interactants, whose nanoBiT signal is independent of Bdf1 BDs.

Since the addition of the C-terminal SmBiT tag had a milder effect on Bdf1 expression levels compared to LgBiT (**Fig. 6b**), we screened Bdf1-SmBiT against LgBiT-tagged partners. Among histones, H2A variant H2A.Z yielded the strongest NanoBiT signal in combination with Bdf1 (**Fig. 6c**). Bdf1-SmBiT was subcloned in an autonomous plasmid under the control of its endogenous promoter and ectopically expressed in *S. cerevisiae*, in a WT or mutant form. Point or deletion mutations that inactivated both Bdf1 BDs yielded a dramatic decrease in signal for assays, using the *BDF1* sequence from either *S. cerevisiae* or *C. glabrata*, strongly confirming specificity of the nanoBiT signal (**Fig. 6d**). Importantly, the nanoBiT signal was significantly lowered by both **24c** and I-BET726, which yielded IC_50_ values of 65 and 2.1 µM, respectively (**Fig. 6e**), comparable to the EC50 values (93 and 3.4 μM) observed in *C. glabrata* growth inhibition assays (**Figs 4g** and **5f**). Inhibition was specific to this protein pair and not observed with other Bdf1 interactants (Swc4) or unrelated nanoBiT assays (Bmh1/Bmh2). These results strongly support Bdf1 BDs as the *in cellulo* target of both **24c** and I-BET726.

### I-BET726 is active against diverse *Candida* species and antifungal-resistant strains

We next profiled the inhibitory activity of I-BET726 against diverse *Candida* species and strains. In HTRF assays with the six Bdf1 BDs from *C. glabrata, C. albicans*, and *C. auris*, I-BET726 showed sub-micromolar activity on four of these BDs (all three BD2 domains plus *Ca*Bdf1 BD1) and low micromolar activity on the other two (**Fig. 7a**). Growth inhibition assays using Bdf1 mutant-expressing *C. glabrata* strains showed that the inhibition observed *in vivo* mirrors the drug susceptibility of the functionally active Bdf1 BD determined *in vitro*: strains expressing Bdf1 mutated in BD1 but with a functionally active BD2 were more susceptible than strains expressing the converse mutants (compare **Fig. 7b** with blue curves in **Fig. 7a**). I-BET726 showed significant efficacy against diverse *Candida* species, especially *C. albicans*, *C. glabrata* and *C. tropicalis* (EC50 values in the low micromolar range), whereas the effect against *C. auris and C. parapsilosis* was more modest and *C. krusei* appeared insensitive (**Fig. 7c**). (Note that the clinical *C. glabrata* isolate tested in **Fig. 7c** was less sensitive than the reference laboratory strain tested in **Fig. 5f**). For the three species for which HTRF data were available, the observed growth inhibitory effect mirrored the potency of I-BET726 against the less sensitive of the two BDs, with EC50 and IC_50_ values related by a factor of approximately 5-7 (compare orange, blue and pink curves in **Fig. 7a** and **c**).

**Figure 7.**
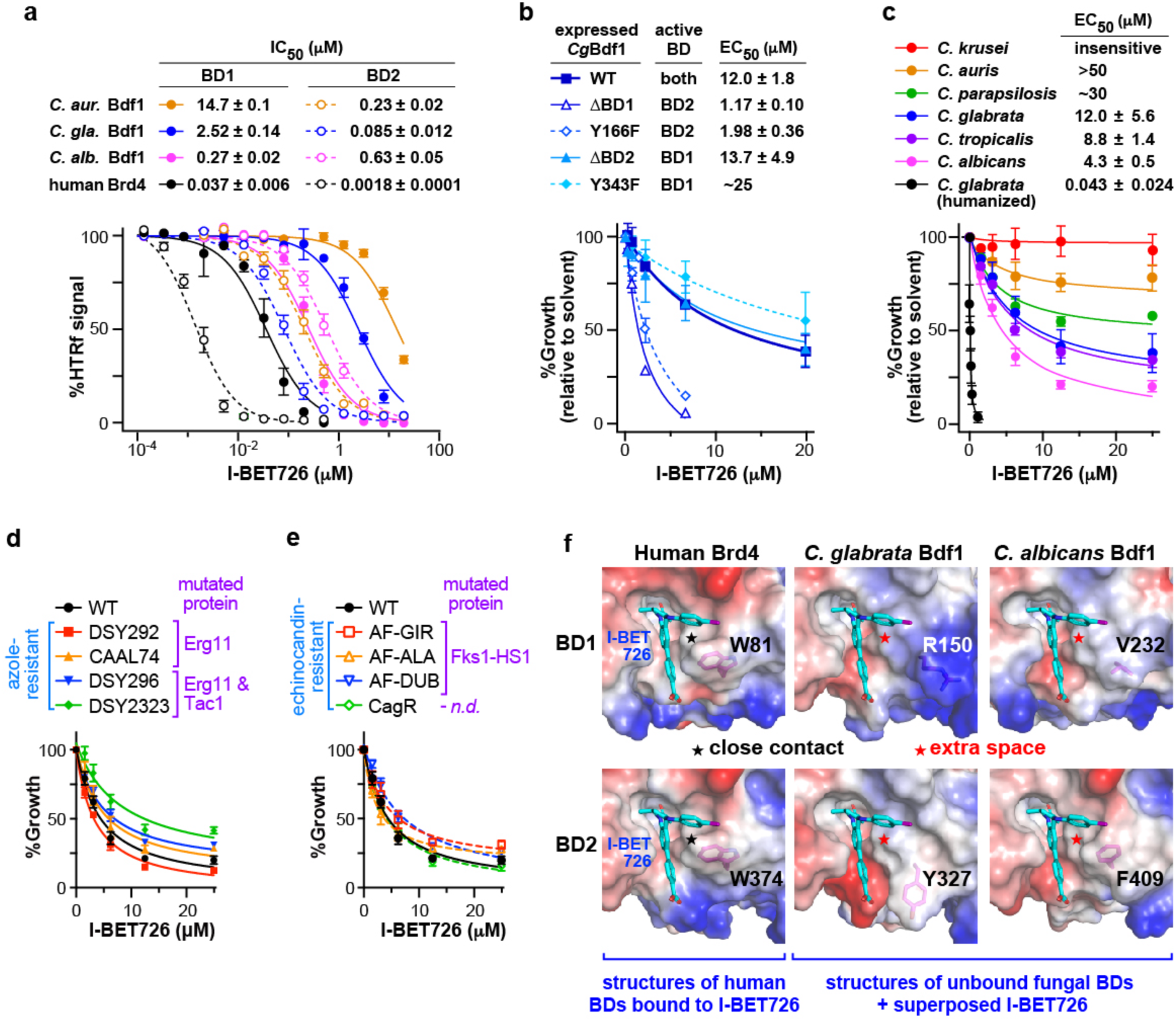
I-BET726 has antifungal activity towards a broad range of *Candida* species. (**a**) HTRF assays showing that I-BET726 inhibits Bdf1 BDs from three phylogenetically diverse *Candida* species, *C. glabrata*, *C. albicans* and *C. auris*. (**b**) *C. glabrata* growth inhibition assays. I-BET726 shows higher efficacy against BD1-inactivated strains compared to BD2-inactivated strains, reflecting its higher potency against BD2 *in vitro*. (**c**) I-BET726 inhibits the growth of several *Candida* species. (**d**, **e**) **I-BET726** inhibits the growth of clinical *C. albicans* strains resistant to (**d**) azoles and (**e**) echinocandins. (**f**) Structural analysis suggests that a modified analog of I-BET726 that clashes with the conserved human Trp in the WPF shelf could confer selectivity for *Candida* Bdf1 BDs.

We next tested the activity of I-BET726 against antifungal-resistant *C. albicans* clinical isolates, including four azole-resistant and four echinocandin-resistant strains (**Table 1** and **Fig. 7d**, **e**). Two of the former group bear mutations in *ERG11*, which encodes the target of azoles (lanosterol 14-*α*-demethylase), while the other two are additionally mutated in *TAC1*, which encodes a transcription factor that regulates the expression of efflux pumps ^44^. Three of the echinocandin-resistant strains bear mutations in *FKS1*, which encodes the enzymatic target of echinocandins (1,3-*β*-glucan synthase), while the fourth has not been genetically characterized. Notably, I-BET726 inhibited the growth of all eight strains with EC50 values (5-10 μM) comparable to that for the reference WT strain. These observations suggest that BET inhibition may be useful for countering the problem of resistance to current antifungal drugs.

**Table 1.** Growth inhibition of C. albicans clinical strains by I-BET726.

| Strains | Resistance | Mutations (ref.<sup>44</sup>) | EC<sub>50</sub> (μM)* |
| --- | --- | --- | --- |
| DSY292 | Azoles | Erg11 (Y132H, G464S, R467K) | 3.07 ± 0.26 |
| CAAL74 | Fluconazole | Erg11 (Y132F, E266D, G448V, V488I) | 5.87 ± 1.03 |
| DSY296 | Azoles | Erg11 (G129A, G464S, G450E)<br>Tac1 (N977D) | 7.36 ± 0.76 |
| DSY2323 | Azoles | Erg11 (S405F)<br>Tac1 (G980E) | 11.5 ± 1.7 |
| AF-GIR | Echinocandins | Fks1-HS1 S645P | 7.13 ± 1.48 |
| AF-ALA | " | Fks1-HS1 S645P | 4.59 ± 0.95 |
| AF-DUB | " | Fks1-HS1 S645P | 6.83 ± 0.26 |
| CagR | " | No characterized | 4.14 ± 0.48 |

\*Data are presented as means ± SD from 3 independent experiments.

Our HTRF assays show that I-BET726 inhibits human BET BDs more potently than the Bdf1 BDs of *C. glabrata* and *C. albicans* (by a factor of 7-70 for BD1 and of 50-350 for BD2; **Fig. 7d**).

Importantly, structural analysis suggests how to invert this selectivity. I-BET726 packs intimately against the WPF Trp residue that is conserved across human BET proteins (**Fig. 7f**). As shown for **24c** (**Fig 4j**), the replacement of this tryptophan by *Cg*Bdf1 residues Arg150 and Tyr327 creates additional space in the binding pocket that may be occupied by an inhibitor with selectivity for the *Cg*Bdf1 BDs. Interestingly, a similar space is available in the binding pocket of *Ca*Bdf1 BDs, where residues Val232 and Phe409 replace the WPF trpytophan (**Fig. 7f**). These observations suggest the potential development of Bdf1-selective analogs of I-BET726 by introducing modifications that clash with the human WPF Trp residue while favourably occupying the extra space available in the *Candida* BD binding pockets.

## Discussion

We previously identified the fungal BET protein Bdf1 as a potential new antifungal target in *C. albicans*^29^. We showed that mutations that inactivate the BDs of Bdf1 are lethal and identified small molecules that inhibited either BD1 or BD2 with high selectivity over human BDs. However, a major obstacle is that the dual inactivation of both Bdf1 BDs is required for lethality, with only a modest effect observed when a single domain is inactivated. Identifying a single compound (or combination of compounds) targeting both BDs with high antifungal potency and selectivity versus human BDs has proved challenging. In addition, to what extent BET inhibition might be useful against other *Candida* species and against drug-resistant strains, and whether a BET inhibitor with broad-spectrum antifungal activity could be envisaged, remained open questions.

To address these, we investigated Bdf1 BD inhibition in *C. glabrata*, the second-most frequent cause of invasive candidiasis, associated with higher overall mortality than *C. albicans* ^7, 31–33, 45^. As in *C. albicans*, Bdf1 is essential in *C. glabrata* and the dual inactivation of both its BDs is required for lethality. Interestingly, the acetylpeptide binding profiles of BD1 and BD2 are more similar for *Cg*Bdf1 than for *Ca*Bdf1, suggesting that dual BD inhibitors might more easily be identified for the former. Indeed, our chemical screen identified several molecules that inhibited *Cg*BD1 and *Cg*BD2 with similar potency. Significantly, minor modifications of a compound originally identified as a *Cg*BD1-selective inhibitor yielded a phenyltriazine derivative, **24c**, that showed low micromolar potency against both *Cg*BDs while maintaining selectivity against human BET BDs.

Our structural analyses of ligand-bound *Cg*BDs greatly elucidates the stereochemical requirements for dual BD activity and fungal-vs-human selectivity. The *Cg*BD2/I-BET151 crystal structure and related structural models reveal that replacing the WPF tryptophan in human BET BDs by an Arg or Tyr residue in *Cg*BD1 and *Cg*BD2, respectively, results in a widened binding pocket predicted to interact less intimately with BET inhibitors I-BET151, JQ1, bromosporine and PFI-1, explaining their decreased potency towards the fungal BDs. Conversely, the *Cg*BD1/**63** crystal structure reveals that **63** achieves high selectivity for *Cg*BD1 by occupying the widened space below the RPF-shelf Arg150 residue, in an orientation that would clash with the Tyr and Trp residues of the *Cg*BD2 YPF and Brd4 WPF shelves. In contrast, the *Cg*BD2/**29** structure and related models show that **29** sits above the Arg, Tyr or Trp residue of the R/Y/WPF shelf, explaining the ability of this compound to inhibit all four *Cg*Bdf1 and Brd4 BDs with similar potency. Interestingly, the structure of *Cg*BD1 bound to **24c**, which selectively inhibits both *Cg*BD1 and *Cg*BD2 over the human BET BDs, reveals a binding orientation that partly overlaps with both those of **29** and **63,** with the **24c** methylfuryl group positioned where the trajectories of these two compounds diverge, immediately next to the first residue in the R/Y/WPF shelf.

Considerable evidence indicates that the antifungal activity of **24c** is an on-target effect. First, **24c** inhibits the growth of the WT strain more effectively than the closely related *Cg*BD1-selective parent compound **24,** which only inhibits strains with BD2-inactivated Bdf1. Second, **24c**, which shows little *in vitro* activity towards human Brd4 BDs, is less effective at inhibiting the growth of strains expressing a humanized variant of Bdf1, in which the two *Cg*BDs have been replaced by their Brd4 counterparts, unlike JQ1 which exhibits the opposite behaviour. Finally, a nanoBiT assay designed to assess the histone binding activity of Bdf1 in yeast cells, reveals that **24c** inhibits this interaction in *S. cerevisiae* with similar potency as observed in *C. glabrata* growth inhibition assays. These results confirm **24c** as the first example of a BET inhibitor that selectively targets fungal BET BDs with antifungal activity against a WT *Candida* strain, establishing an important proof of concept. Furthermore, the small size (294 Da) and relatively simple structure of **24c** compared to that of most currently investigated BET inhibitors (typically 350-500 Da) suggest that the antifungal potency of this compound may be significantly improved through chemical optimization.

Our use of *C. glabrata* strains expressing humanized Bdf1 led us to discover the remarkable potency of BET inhibitor I-BET726 against the humanized strain, with an EC50 value in the low nanomolar range. This observation strongly supports the feasibility of developing Bdf1 BD inhibitors as highly potent antifungal agents. Interestingly, I-BET726 showed significant growth inhibitory activity against a broad range of unmodified *Candida* species and strains, including azole- and echinocandin-resistant clinical isolates, suggesting that BET inhibition may provide a useful strategy to combat antifungal resistance. Although I-BET726 has higher potency for human than for fungal BET BDs, our structural analyses of *Cg*BD-selective compounds suggest potential modification strategies that could invert this selectivity.

Taken together, the above findings establish Bdf1 as a potential new antifungal target in *C. glabrata* and strongly support the development of BET inhibitors as a novel class of antifungal therapeutics with the potential for broad spectrum anti-*Candida* activity.

## METHODS

### Chemicals

BET inhibitors I-BET151, bromosporine, PFI-1 and I-BET762 were purchased from Sigma. I-BET726, RVX208 and ABBV744 were purchased from Euromedex while JQ1 was from Clinisciences (Nanterre, France). Screening validation and derived compounds were purchased from ChemDiv as powder and dissolved in dimethylsulfoxide (DMSO) without further purification. Compounds **24k**, **24l** and **24m** were synthesized in-house.

### Protein expression and purification

#### Proteins used for crystallization

DNA fragments encoding *Cg*Bdf1 BD1 (residues 128–237) or BD2 (residues 304–418) were PCR amplified from genomic DNA and cloned into a pETM11 vector as fusion constructs bearing an N-terminal His tag followed by a tobacco etch virus (TEV) protease cleavage site. Transformed *E. coli* strains BL21(DE3) (New England Biolabs, ref. C2527I) were grown in LB medium containing kanamycin (50 µg/mL) at 37 ^°^C until an OD600 of 0.5, induced with 1 mM IPTG and further incubated at 18 ^°^C for 12–20 h before collecting. Cells were resuspended in buffer A (50 mM Tris-HCl pH 7.5, 300 mM NaCl, 10 % glycerol, 25 mM imidazole, 5 mM *β*-mercaptoethanol and protease inhibitors) and lysed by sonication. The cleared lysate was incubated with Ni-NTA resin (Qiagen) and washed with buffer A containing 0.5 M NaCl. Proteins were eluted with 250 mM imidazole, dialysed overnight in the presence of His-tagged TEV protease against buffer A (containing no imidazole). After dialysis, Ni-NTA resin was used to remove His-tagged species. Proteins were further purified on a Superdex 75 10/300 (GE Healthcare) column in 50 mM Hepes pH 7.5, 150 mM NaCl, 0.5 mM dithiothreitol (DTT). Proteins were concentrated to >20 mg/mL on a Centricon device (Millipore).

#### Proteins used for HTRF assays

GST-tagged human Brd4 BD2 (residues 349–460) was purchased from Reaction Biology Corp. Human Brd4 BD1 (residues 22–204), CaBdf1 BD1 (residues 193–327) and BD2 (residues 361–501), CgBdf1 BD1 (residues 120–248) and BD2 (residues 289–411) and CauBdf1 BD1 (residues 127–257) and BD2 (residues 287–416) were cloned into a pGEX4t1 vector as GST-tagged fusion proteins. Expression in *E. coli* strain BL21(DE3) cells was performed as for His-tagged constructs. Collected cells resuspended in 50 mM Tris-HCl pH 7.5, 150 mM NaCl and protease inhibitors were lysed by sonication. The clarified lysate was incubated with glutathione sepharose (GE Healthcare) and then washed with 50 mM Tris-HCl pH 7.5, 500 mM NaCl and 1 % NP-40. Proteins were eluted with 10 mM glutathione and further purified on a Superdex 200 10/300 (GE Healthcare) column in 50 mM Hepes pH 7.5, 150 mM NaCl, 0.5 mM DTT. Glycerol was added additionally to a final concentration of 30%.

### Pull-down assay

Pull-down assays were performed as described^29^. Biotinylated peptides corresponding to non-acetylated and tetra-acetylated (H4K5acK8acK12acK16ac) histone H4 tails were synthesized by Covalab (Villeurbanne, France) and immobilized on Streptavidin-coated magnetic beads (Dynabeads MyOne Streptavidin C1; Thermo Fisher) according to the manufacturer’s instructions. Beads were incubated with 1.25 mg of GST-tagged *Cg*Bdf1 BD1 or BD2 in binding buffer (50 mM Tris pH 7.4, 150 mM NaCl, 0.1 % NP-40, 10 % glycerol 10 %, 1 mM DTT) in a volume of 250 mL for 2h at 4 °C and subsequently washed in binding buffer containing 500 mM NaCl. Bound proteins were eluted by boiling in SDS–PAGE sample loading buffer and analyzed by western blot using an anti-GST antibody (GE Healthcare).

### Histone peptide array assay

MODified Histone Peptide Array Kit was purchased from Active motif (Ref. 13005). Array was prepared as described in the manufacturer’s instructions. 1 µM of proteins in binding buffer (50 mM Tris pH 7.4, 150 mM NaCl, 0.1 %, NP-40, 10 % glycerol, 1 mM DTT) were incubated with the array at 4 °C for two hours. The array was then washed three times with binding buffer. Anti-GST antibody (GE Healthcare) was used for the detection of GST-tagged bromodomains.

### HTRF assay

HTRF reagents and buffers were purchased from Cisbio Bioassays. The assay used a terbium (III) cryptate donor reagent conjugated to an anti-GST antibody (MAb anti-GST-Tb; ref. 61GSTTLA), a streptavidin-conjugated acceptor reagent (streptavidin-d2; ref. 610SADLA) and Cisbio proprietary buffers (EPIgeneous Binding Domain Diluent and Detection buffer; refs. 62DLBDDF and 62DB2FDG, respectively). Incubation with GST-tagged BDs and biotinylated H4ac4 brings the donor and acceptor into close proximity and allows FRET. The non-biotinylated H4ac4 peptide competes for binding and was used as a positive control of inhibition. GST-tagged proteins in 25 mM Hepes pH 7.5, 150 mM NaCl, 0.5 mM DTT were assayed at a final concentration of 5 nM. Biotinylated H4ac4 peptides were used at a final concentration of 50, 600, 300, 400, 30, 250, 176 or 1,450 nM in assays involving Brd4 BD1, Brd4 BD2, CaBdf1 BD1, CaBdf1 BD2, CgBdf1 BD1, CgBdf1 BD2, CauBdf1 BD1 and CauBdf1 BD2, respectively. The antibody-conjugated donor was used at 0.5 nM and the streptavidin-conjugated acceptor was used at 1/8 of the H4ac4 peptide concentration. Inhibitors were tested by performing an eleven-point 2.5-fold dilution series with a maximal final concentration of 20 mM. These concentrations allowed a fixed DMSO concentration at 0.2 %, critical for a Z’ factor ≥ 0.8. Components were incubated at 4 ^°^C for 4 h (all BD1) or for 24 h (all BD2). Experiments were performed in 384-well white plates (Greiner ref. 781080) in a volume of 16 μL and analyzed in a ClarioStar plate reader (BMG LABTECH, excitation at 330 nm and emission at 620 and 665 nm, corresponding to the donor and acceptor emission peaks, respectively; the 665/620 ratio is used to calculate the specific HTRF signal) with an integration delay of 60 µs and an integration time of 400 µs.

### High-throughput chemical screening

The HTRF assay described above was miniaturized in a 5 µL volume in 1,536-well black plates. Approximately 100,000 compounds comprising the soluble diversity (ChemDiv), targeted diversity (ChemDiv) and 30K diversity (LifeChem) collections were dispensed into wells by an Echo acoustic liquid dispenser. A master mix comprising MAb anti-GST-Tb donor, streptavidin-d2 acceptor, GST-tagged Cg Bdf1 BD1 protein, biotinylated H4ac4 peptide was then added and the plates incubated for 4 h prior to reading. The primary screen was performed with compounds at a final concentration of 20 µM, corresponding to a final DMSO concentration of 1 %. Hits were initially confirmed by repeating the assay at a single concentration in triplicate, and subsequently by dose–response curves constructed using eight-point dilutions between 0 and 20 µM.

### Crystallization and crystal structure determination

Initial crystallization conditions were identified by the sitting drop vapor diffusion method at 4 °C using a Cartesian PixSys 4200 crystallization robot at the High Throughput Crystallization Laboratory of the EMBL Grenoble Outstation (https://htxlab.embl.fr). Crystals used for data collection were obtained by the hanging drop method at 4 °C by mixing 1 μL of the protein or protein/inhibitor sample with 1 μL of the reservoir solution, as follows. Crystals were harvested directly from the crystallization drop unless otherwise specified. Unbound *Cg*Bdf1 BD1 (25 mg/mL) was mixed with 25% (w/v) PEG 1500 and 0.1 MIB (malonic acid:imidazole:boric acid in 2:3:3 molar ratio) buffer (pH 9). Unbound CgBdf1 BD2 (40 mg/mL) was mixed with 0.1 M CH3COONa (pH 4.6) and 33 % (v/v) MPD. CgBdf1 BD2 bound to I-BET151 was crystallized by mixing a solution of 15 mg/mL protein and 2.2 mM inhibitor with 0.9 M (NH4)2SO4 and 0.1 M sodium cacodylate (pH 6.5); crystals were harvested in a cryoprotectant containing 1.1 M (NH4)2SO4, 0.1 M sodium cacodylate (pH 6.5) and 30% (v/v) glycerol. CgBdf1 BD1 bound to Compound **24c** was crystallized by mixing a solution of 35 mg/mL protein and 5.4 mM inhibitor with 0.072 M NaH2PO4 and 1.73 M K2HPO4 (pH 8.2). Crystals were flash-cooled in liquid nitrogen.

Diffraction data collected at beamlines of the ESRF and Soleil were processed using XDS^46^ and programs of the CCP4 suite^47^. Structures were solved by molecular replacement using Phaser^48^, improved by manual building using Coot^49^, and refined with Phenix^50^. Data collection and refinement statistics are summarized in **Suppl. Table 1**.

### Generation of *C. albicans* mutant strains

Plasmids used in this study are listed in **Suppl. Table 5**. All DNA fragments were fused in pCR2.1-TOPO (genome integration cassette) or pGRB2.0 (rescue plasmids) vectors using a Gibson assembly kit (New England Biolabs) and validated by sequencing. The *BDF1* point mutant plasmids were obtained using the QuikChange II Site-directed mutagenesis kit (Agilent) with the *BDF1* plasmid pJG267. *C. glabrata* was transformed by a classic lithium acetate-based procedure, as previously described^51^ with minor changes (incubation at 30 °C for 30 min instead of overnight). The pMET3 promoter sequence was recycled from the plasmid pFA-ARG4-MET3p ^52^. *C. glabrata* strains used in this study are listed in **Suppl. Table 6**.

### Growth assays

Growth assays were performed as described^29^ with minor changes.

#### Growth on solid media

*C. glabrata* strains were grown in SC or SC-U (SC medium without uracil) media to an OD600 of 0.5–0.8, pelleted and resuspended in sterile water at a final OD600 of 0.13. Cells were spotted on solid media in a threefold dilution series starting at an OD600 of 0.13. Plates were incubated at 30 ^°^C for 1 day before imaging.

#### Growth on liquid media with C. glabrata lab strains

Cells were grown at 30 °C in SC+M+C-U medium (SC medium with 5 mM methionine, 0.25 mM cysteine, without uracil) to an OD600 of 0.5–0.8. For the evaluation of growth defects related to Bdf1 mutations, log-phase growing cells were counted using a Neubauer chamber and diluted in liquid media to a final concentration of 13,500 cells per mL (67,500 cells per mL for *BDF1*-hBD-FLAG and *BDF1*-hBD1-FLAG pMET strains) per well in a sterile 96-well plate. Plates were incubated at 30 ^°^C and OD600 was measured using a Multiskan FC Microplate Photometer (Thermo Fisher). For the evaluation of chemical compounds, *C. glabrata* pMET strains were grown in SC+M+C-U medium 24 h before counting and then seeded in 96-well plates at the same concentration as mentioned above, with or without chemical compounds.

#### Growth of clinical strains in liquid media

Strains are presented in **Suppl. Table**. Cells from a single colony were diluted in 0.85 % NaCl before being counted using a Neubauer chamber. They were diluted to a final concentration of 6,500 cells per mL per well for all clinical strains of *C. albicans* and *C. glabrata*; 65,000 cells per mL per well for *C. auris*; 13,500 cells per mL per well for *C. parapsilosis*; 3,375 cells per mL per well for *C. krusei* and 1,687.5 cells per mL per well for *C. tropicalis*, in a sterile 96-well plate. Cells were then grown in SC medium with or without chemical compounds. The significance of growth defects were assessed using Holm-Sidak method by GraphPad Prime 7.

### Analysis of whole-cell extracts and antibodies

*C. glabrata* strains were grown at 30°C in liquid media to an OD600 of 0.5–0.8. Cells were lysed in FastprepTM (MPBiologicals) twice at 6 m/s for 30 s with intermediate incubation on ice. FLAG antibody was purchased from Sigma-Aldrich (ref. F3165) and endogenous CgBdf1 detected with an antibody developed internally against Bdf1 from *S. cerevisiae* ^27^.

### Cytotoxicity assays on human cells

Cytotoxicity assays were performed as described^29^ (with minor changes. Proliferation of human cells was assessed using an MTT colorimetric assay (Cell Proliferation Kit I, Roche). HeLa (epithelial cells, ATCC number CCL-2) and IMR90 (primary fibroblasts cells, ATCC number CCL-186) cells were cultured in humidified atmosphere (37 ^°^C and 5 % CO2) in DMEM medium containing 10 % heat inactivated fetal calf serum and 2 mM glutamine. Cells were seeded at a concentration of 5,000 (HeLa) or 5,600 (IMR90 cells) per well in 100 µL culture medium containing the test compound (Compound 3-3, amphotericin B or fluconazole) into 96 wells microplates (Falcon ref. 353072). Plates were incubated at 37 ^°^C and 5 % CO2 for 24 h before adding 10 mL of MTT labelling reagent (final concentration 0.5 mg/mL) to each well. After incubating for a further 4 h, 100 m-L of the solubilization solution were added in each well. Plates were allowed to stand overnight in the incubator before measuring the absorbance at 570 nm and at 690 nm in a ClarioStar plate reader. The values of A570 nm–A690 nm were normalized relative to that obtained with vehicle (0.2 % DMSO, 0.8 % ethanol) and plotted against compound concentration.

## SUPPLEMENTARY INFORMATION

**Supplementary Table 1.**
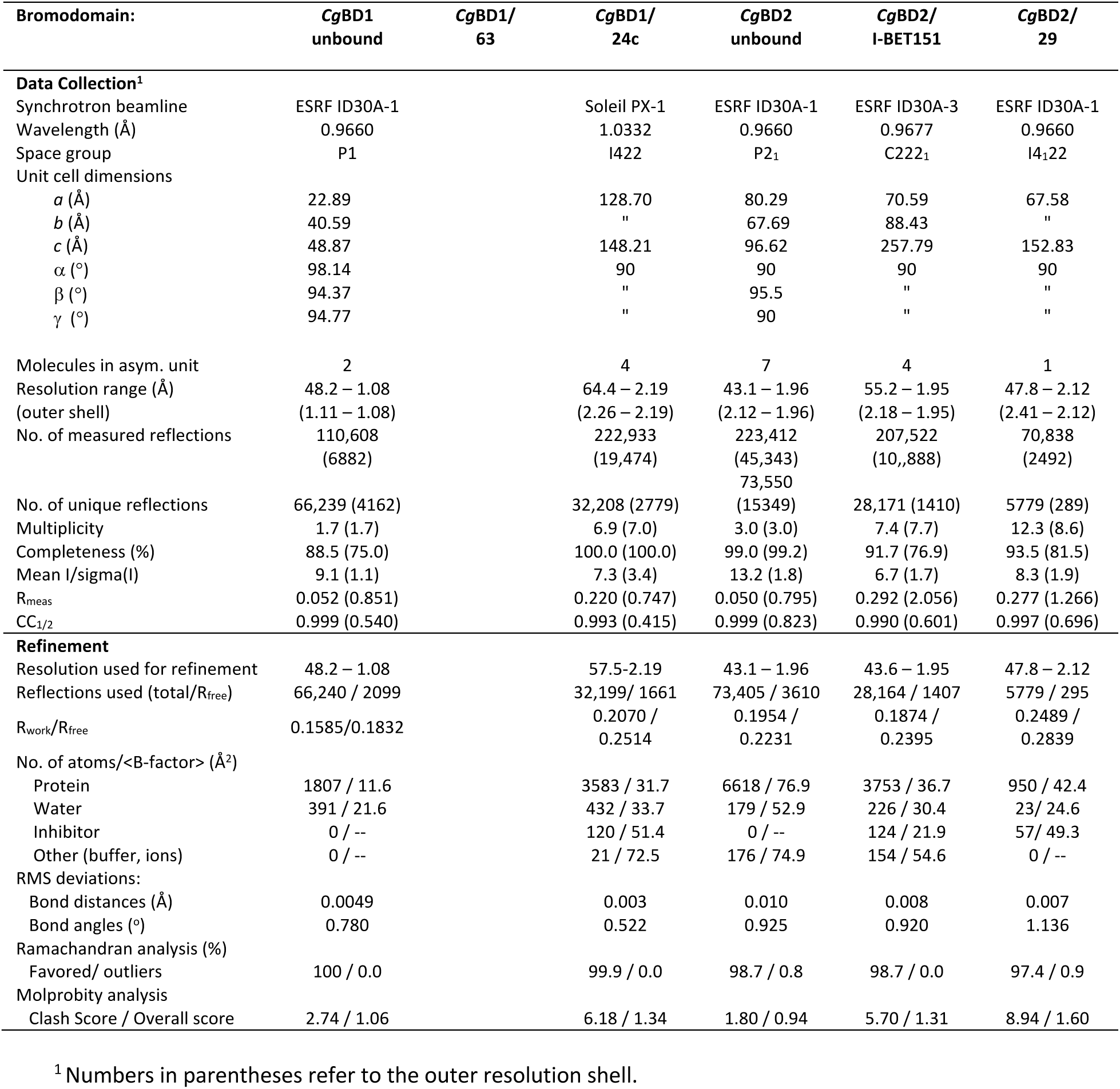
Crystallographic data collection and refinement statistics.

**Supplementary Table 2.**
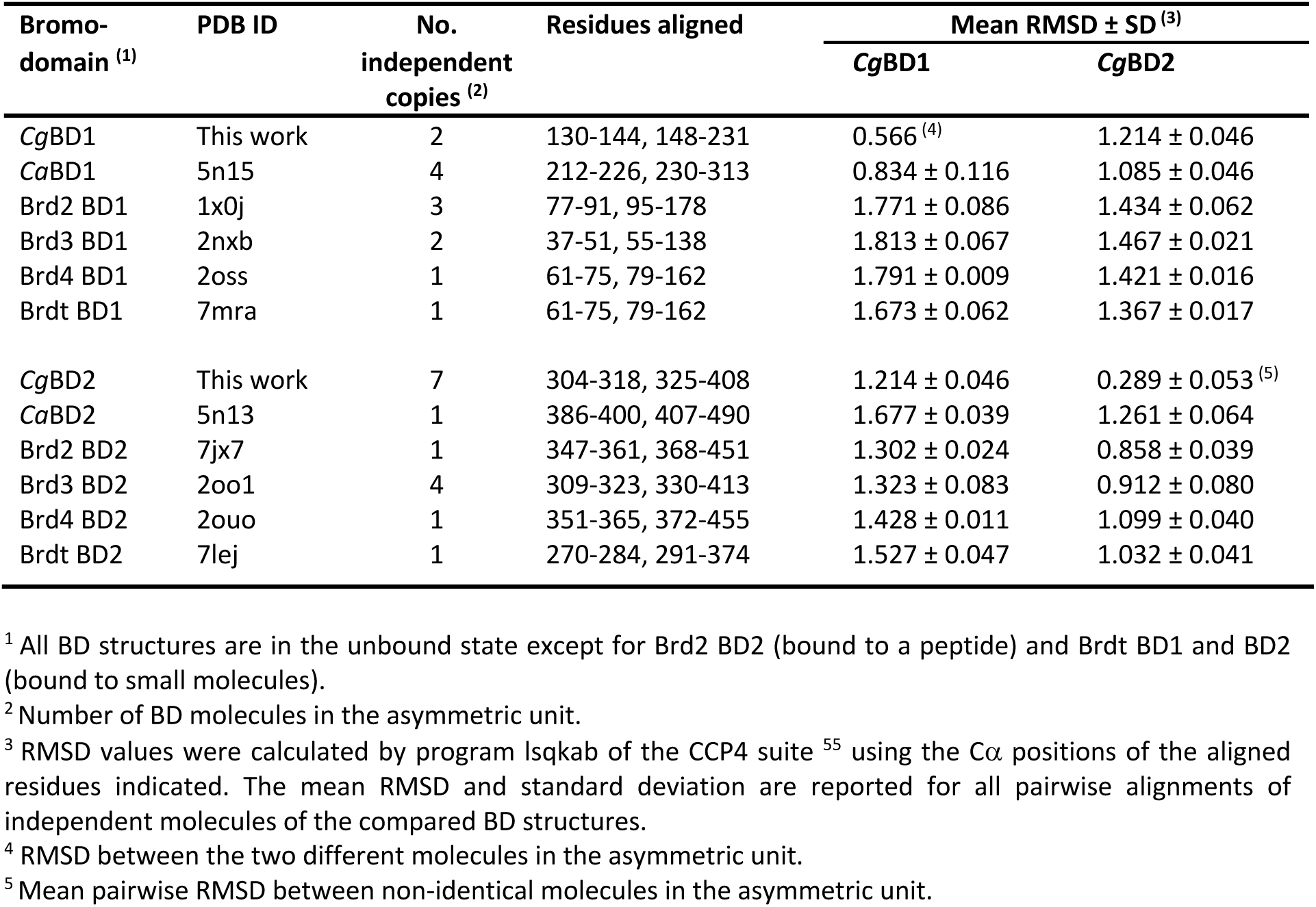
RMSD values for alignments of *Cg*Bdf1 BDs with *Ca*Bdf1 and human BET BDs.

| Bromo-domain <sup>(1)</sup> | PDB ID | No. independent copies <sup>(2)</sup> | Residues aligned | Mean RMSD ± SD <sup>(3)</sup> | |
| --- | --- | --- | --- | --- | --- |
|  |  |  |  | *CgBD1* | *CgBD2* |
| *CgBD1* | This work | 2 | 130-144, 148-231 | 0.566 <sup>(4)</sup> | 1.214 ± 0.046 |
| *CaBD1* | 5n15 | 4 | 212-226, 230-313 | 0.834 ± 0.116 | 1.085 ± 0.046 |
| Brd2 BD1 | 1x0j | 3 | 77-91, 95-178 | 1.771 ± 0.086 | 1.434 ± 0.062 |
| Brd3 BD1 | 2nxb | 2 | 37-51, 55-138 | 1.813 ± 0.067 | 1.467 ± 0.021 |
| Brd4 BD1 | 2oss | 1 | 61-75, 79-162 | 1.791 ± 0.009 | 1.421 ± 0.016 |
| Brdt BD1 | 7mra | 1 | 61-75, 79-162 | 1.673 ± 0.062 | 1.367 ± 0.017 |
| *CgBD2* | This work | 7 | 304-318, 325-408 | 1.214 ± 0.046 | 0.289 ± 0.053 <sup>(5)</sup> |
| *CaBD2* | 5n13 | 1 | 386-400, 407-490 | 1.677 ± 0.039 | 1.261 ± 0.064 |
| Brd2 BD2 | 7jx7 | 1 | 347-361, 368-451 | 1.302 ± 0.024 | 0.858 ± 0.039 |
| Brd3 BD2 | 2oo1 | 4 | 309-323, 330-413 | 1.323 ± 0.083 | 0.912 ± 0.080 |
| Brd4 BD2 | 2ouo | 1 | 351-365, 372-455 | 1.428 ± 0.011 | 1.099 ± 0.040 |
| Brdt BD2 | 7lej | 1 | 270-284, 291-374 | 1.527 ± 0.047 | 1.032 ± 0.041 |

<sup>1</sup> All BD structures are in the unbound state except for Brd2 BD2 (bound to a peptide) and Brdt BD1 and BD2 (bound to small molecules).
<sup>2</sup> Number of BD molecules in the asymmetric unit.
<sup>3</sup> RMSD values were calculated by program lsqkab of the CCP4 suite <sup>55</sup> using the C $\alpha$ positions of the aligned residues indicated. The mean RMSD and standard deviation are reported for all pairwise alignments of independent molecules of the compared BD structures.
<sup>4</sup> RMSD between the two different molecules in the asymmetric unit.
<sup>5</sup> Mean pairwise RMSD between non-identical molecules in the asymmetric unit.

**Supplementary Table 3.** IC_50_ estimates for HTS hit compounds in preliminary HTRF assay.

| Com-<br>pound | IC<sub>50</sub> (μM) <sup>(1)</sup> |  |  |  | Com-<br>pound | IC<sub>50</sub> (μM) |  |  |  |
| --- | --- | --- | --- | --- | --- | --- | --- | --- | --- |
|  | CgBdf1<br>BD1 | CgBdf1<br>BD2 | Brd4<br>BD1 | Brd4<br>BD2 |  | CgBdf1<br>BD1 | CgBdf1<br>BD2 | Brd4<br>BD1 | Brd4<br>BD2 |
| **1** | 9.4 | >20<sup>(2)</sup> | >20 | >20 | **33** | 6.1 | *L* | *L* | *L* |
| **2\*** <sup>(3)</sup> | 4.4 | 1.1 | 13 | 2.8 | **34** | >20 | >20 | *L* | *L* |
| **3** | 19 | >20 | >20 | >20 | **35** | >20 | *L* | >20 | *L* |
| **4** | >20 | *L*<sup>(2)</sup> | *L* | *L* | **36** | *L* | *L* | *L* | >20 |
| **5\*** | 1.8 | 1.0 | 2.5 | 1.4 | **37** | >20 | *L* | *L* | *L* |
| **6** | 19 | 8.2 | >20 | 2.7 | **38** | >20 | 6.5 | *L* | >20 |
| **7** | *L* | *L* | >20 | >20 | **39** | 6.7 | *L* | *L* | *L* |
| **8** | 6.4 | 6.3 | 18 | 12 | **40** | >20 | *L* | *L* | >20 |
| **9** | *L* | *L* | *L* | *L* | **41** | >20 | >20 | *L* | >20 |
| **10** | 5.0 | >20 | *L* | *L* | **42** | 5.9 | 8.8 | 19 | 14 |
| **11** | >20 | *L* | *L* | *L* | **43\*** | 2.7 | 13 | >20 | >20 |
| **12** | >20 | *L* | >20 | 10 | **44** | 19 | >20 | >20 | *L* |
| **13** | >20 | *L* | *L* | *L* | **45\*** | 4.2 | *L* | >20 | *L* |
| **14** | *L* | *L* | >20 | >20 | **46** | >20 | *L* | *L* | *L* |
| **15** | *L* | *L* | *L* | >20 | **47** | 11 | *L* | >20 | *L* |
| **16\*** | 1.8 | 0.30 | 1.2 | 1.5 | **48** | >20 | *L* | *L* | >20 |
| **17** | 7.0 | 21 | 9.5 | >20 | **49** | >20 | *L* | *L* | *L* |
| **18** | 7.8 | 13 | 16 | >20 | **50** | >20 | *L* | *L* | *L* |
| **19** | >20 | *L* | >20 | *L* | **51** | 12 | >20 | 12 | >20 |
| **20** | 23 | >20 | >20 | >20 | **52** | >20 | *L* | *L* | *L* |
| **21** | >20 | >20 | >20 | 3.5 | **53** | >20 | *L* | *L* | *L* |
| **22** | 21 | >20 | >20 | >20 | **54** | >20 | *L* | *L* | *L* |
| **23** | >20 | *L* | *L* | *L* | **55\*** | 0.11 | *L* | >20 | *L* |
| **24\*** | 2.5 | >20 | >20 | >20 | **56\*** | 0.36 | *L* | *L* | *L* |
| **25** | >20 | >20 | >20 | >20 | **57** | >20 | *L* | *L* | *L* |
| **26** | 8.4 | *L* | >20 | *L* | **58** | >20 | >20 | >20 | >20 |
| **27** | 5.9 | 17 | >20 | *L* | **59** | 14 | *L* | >20 | >20 |
| **28** | >20 | >20 | >20 | 3.5 | **60** | >20 | >20 | >20 | >20 |
| **29\*** | 1.1 | 2.3 | 2.9 | 0.44 | **61** | >20 | >20 | >20 | *L* |
| **30\*** | 0.64 | *L* | >20 | >20 | **62** | >20 | *L* | >20 | *L* |
| **31** | 6.3 | *L* | >20 | *L* | **63\*** | 0.62 | *L* | >20 | *L* |
| **32\*** | 2.9 | >20 | >20 | >20 |  |  |  |  |  |

<sup>1</sup> All IC<sub>50</sub> values were determined from a single-replicate HTRF assay.
<sup>2</sup> IC<sub>50</sub> values for compounds yielding weak (<40%) or no detectable inhibition at the maximal tested concentration are indicated as >20 and *L* (large), respectively.
<sup>3</sup> Compounds retained for further characterization are indicated by an asterisk.

**Supplementary Table 4.** IC_50_ values of HTS hits with highest potency towards *Cg*BD1.

| Compound | IC<sub>50</sub> (μM) <sup>(1)</sup> |  |  |  |
| --- | --- | --- | --- | --- |
|  | CgBD1 | CgBD2 | hBrd4 BD1 | hBrd4 BD2 |
| **2** | 4.97 ± 0.77 | 0.97 ± 0.25 | 21.8 ± 3.6 | 2.93 ± 0.57 |
| **5** | 1.95 ± 0.35 | 2.22 ± 1.68 | 3.07 ± 0.52 | 1.33 ± 0.07 |
| **16** | 4.65 ± 2.74 | 0.23 ± 0.08 | 1.13 ± 0.29 | 1.57 ± 0.10 |
| **24** | 2.23 ± 0.26 | >20 <sup>(2)</sup> | >20 | >20 |
| **29** | 1.81 ± 1.14 | 2.06 ± 0.23 | 2.73 ± 0.34 | 0.52 ± 0.20 |
| **30** | 0.96 ± 0.32 | *L* <sup>(2)</sup> | >20 | >20 |
| **32** | 2.84 ± 0.59 | >20 | >20 | >20 |
| **43** | 3.20 ± 0.59 | *n.d.* <sup>(3)</sup> | >20 | >20 |
| **45** | 3.08 ± 1.86 | *L* | >20 | *L* |
| **55** | 0.10 ± 0.03 | *L* | *L* | *L* |
| **56** | 0.38 ± 0.17 | *L* | *L* | *L* |
| **63** | 0.61 ± 0.10 | *L* | 25.9 ± 7.9 | *L* |

<sup>1</sup> Data are presented as means ± SD for *n*=3 independent experiments.
<sup>2</sup> IC<sub>50</sub> values for compounds yielding weak (<40%) or no detectable inhibition at the maximal tested concentration (20 μM) are indicated as >20 and *L* (large), respectively.
<sup>3</sup> A reliable value could not be determined because of unusually large scatter of the data.

**Supplementary Table 5.**
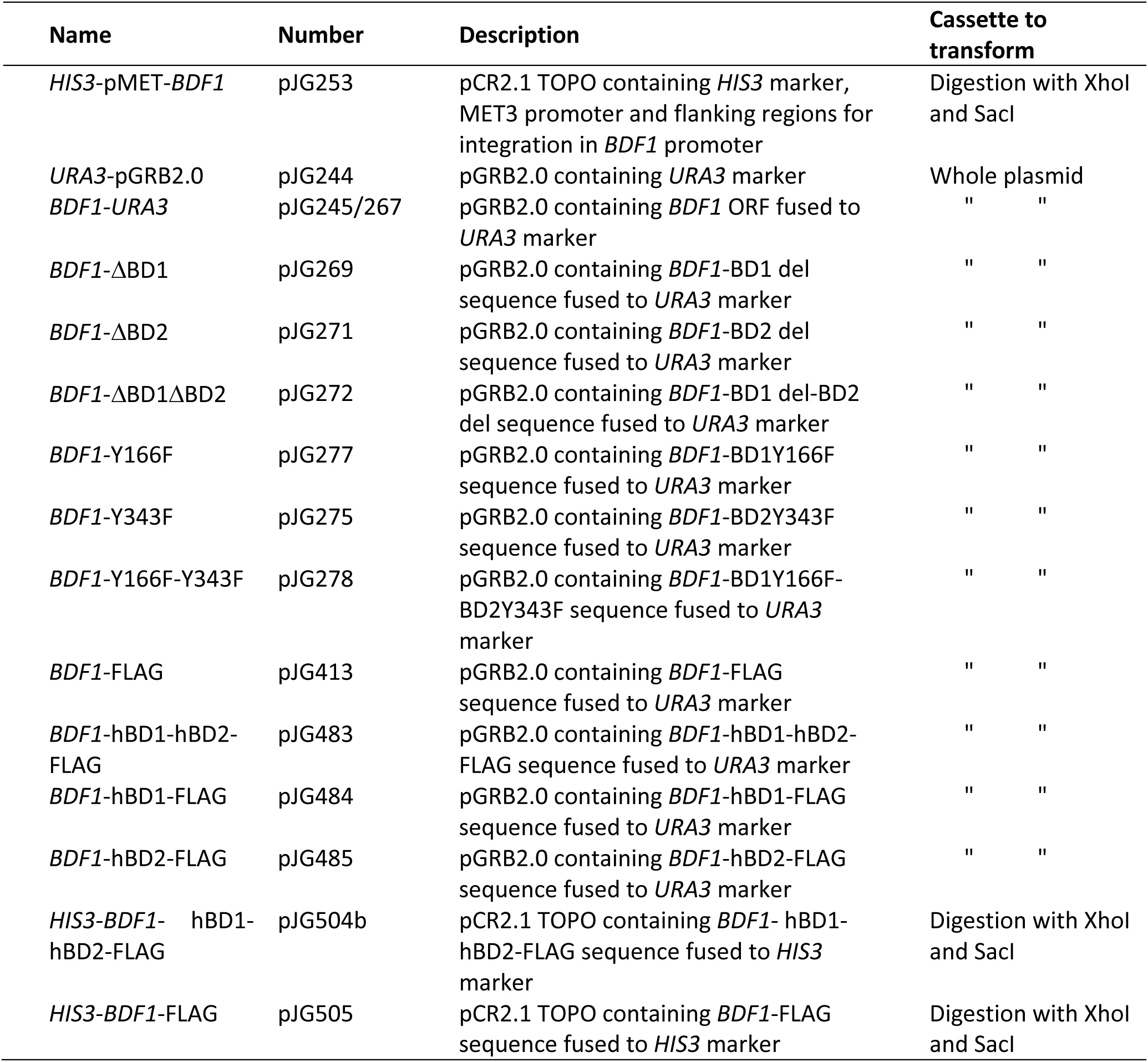
Plasmids used in this study.

| **Name** | **Number** | **Description** | **Cassette to transform** |
| --- | --- | --- | --- |
| *HIS3*-pMET-*BDF1* | pJG253 | pCR2.1 TOPO containing *HIS3* marker, MET3 promoter and flanking regions for integration in *BDF1* promoter | Digestion with XhoI and SacI |
| *URA3*-pGRB2.0 | pJG244 | pGRB2.0 containing *URA3* marker | Whole plasmid |
| *BDF1-URA3* | pJG245/267 | pGRB2.0 containing *BDF1* ORF fused to *URA3* marker | " " |
| *BDF1*-ΔBD1 | pJG269 | pGRB2.0 containing *BDF1*-BD1 del sequence fused to *URA3* marker | " " |
| *BDF1*-ΔBD2 | pJG271 | pGRB2.0 containing *BDF1*-BD2 del sequence fused to *URA3* marker | " " |
| *BDF1*-ΔBD1ΔBD2 | pJG272 | pGRB2.0 containing *BDF1*-BD1 del-BD2 del sequence fused to *URA3* marker | " " |
| *BDF1*-Y166F | pJG277 | pGRB2.0 containing *BDF1*-BD1Y166F sequence fused to *URA3* marker | " " |
| *BDF1*-Y343F | pJG275 | pGRB2.0 containing *BDF1*-BD2Y343F sequence fused to *URA3* marker | " " |
| *BDF1*-Y166F-Y343F | pJG278 | pGRB2.0 containing *BDF1*-BD1Y166F-BD2Y343F sequence fused to *URA3* marker | " " |
| *BDF1*-FLAG | pJG413 | pGRB2.0 containing *BDF1*-FLAG sequence fused to *URA3* marker | " " |
| *BDF1*-hBD1-hBD2-FLAG | pJG483 | pGRB2.0 containing *BDF1*-hBD1-hBD2-FLAG sequence fused to *URA3* marker | " " |
| *BDF1*-hBD1-FLAG | pJG484 | pGRB2.0 containing *BDF1*-hBD1-FLAG sequence fused to *URA3* marker | " " |
| *BDF1*-hBD2-FLAG | pJG485 | pGRB2.0 containing *BDF1*-hBD2-FLAG sequence fused to *URA3* marker | " " |
| *HIS3*-*BDF1*-hBD1-hBD2-FLAG | pJG504b | pCR2.1 TOPO containing *BDF1*-hBD1-hBD2-FLAG sequence fused to *HIS3* marker | Digestion with XhoI and SacI |
| *HIS3*-*BDF1*-FLAG | pJG505 | pCR2.1 TOPO containing *BDF1*-FLAG sequence fused to *HIS3* marker | Digestion with XhoI and SacI |

**Supplementary Table 6.** *C. glabrata* strains used in this study.

| Name | Number | Parent | Genotype | Containing plasmid |
| --- | --- | --- | --- | --- |
| ATCC2001 | Clin02 | - | Wildtype | - |
| ATCC2001 HTU (ATCC200989) | CgJG1 | - | *his3Δ trp1Δ ura3Δ* | - |
| ATCC2001 HTL | CgJG72 | - | *leu2::trp1::his3* | - |
| WT *BDF1* + Empty plasmid | CgJG26 | CgJG1 | *ura3::trp1::his3* | pJG244 |
| pMET-*BDF1* | CgJG12 | CgJG1 | pMET-*BDF1::HIS3 his3Δ trp1Δ ura3Δ* | - |
| pMET-*BDF1* + WT *BDF1* | CgJG19/20 | CgJG12 | pMET-*BDF1::HIS3 his3Δ trp1Δ ura3Δ* | pJG245 |
| pMET-*BDF1* + *BDF1*-ΔBD1 | CgJG35/36 | CgJG12 | pMET-*BDF1::HIS3 his3Δ trp1Δ ura3Δ* | pJG269 |
| pMET-*BDF1* + *BDF1*-ΔBD2 | CgJG39/40 | CgJG12 | pMET-*BDF1::HIS3 his3Δ trp1Δ ura3Δ* | pJG271 |
| pMET-*BDF1* + *BDF1*-ΔBD1ΔBD2 | CgJG43/44 | CgJG12 | pMET-*BDF1::HIS3 his3Δ trp1Δ ura3Δ* | pJG272 |
| pMET-*BDF1* + *BDF1*-BD1 Y-F | CgJG1/52 | CgJG12 | pMET-*BDF1::HIS3 his3Δ trp1Δ ura3Δ* | pJG277 |
| pMET-*BDF1* + *BDF1*-BD2 Y-F | CgJG47/49 | CgJG12 | pMET-*BDF1::HIS3 his3Δ trp1Δ ura3Δ* | pJG275 |
| pMET-*BDF1* + *BDF1*-BD1&BD2 Y-F | CgJG55/56 | CgJG12 | pMET-*BDF1::HIS3 his3Δ trp1Δ ura3Δ* | pJG278 |
| pMET-*BDF1* + *BDF1*-FLAG | CgJG80 | CgJG12 | pMET-*BDF1::HIS3 his3Δ trp1Δ ura3Δ* | pJG413 |
| pMET-*BDF1* + *BDF1*-hBD-FLAG | CgJG88 | CgJG12 | pMET-*BDF1::HIS3 his3Δ trp1Δ ura3Δ* | pJG483 |
| pMET-*BDF1* + *BDF1*-hBD1-FLAG | CgJG89 | CgJG12 | pMET-*BDF1::HIS3 his3Δ trp1Δ ura3Δ* | pJG484 |
| pMET-*BDF1* + *BDF1*-hBD2-FLAG | CgJG90 | CgJG12 | pMET-*BDF1::HIS3 his3Δ trp1Δ ura3Δ* | pJG485 |
| *BDF1*-hBD | CgJG92 | CgJG72 | *BDF1-hBD\*::HIS3 his3Δ trp1Δ ura3Δ* | - |
| *BDF1*-FLAG | CgJG95b | CgJG72 | *BDF1-FLAG::HIS3 his3Δ trp1Δ ura3Δ* | - |
| *BDF1*-hBD-FLAG | CgJG93b | CgJG72 | *BDF1-hBD\*-FLAG::HIS3 his3Δ trp1Δ ura3Δ* | - |

**Supplementary Table 7.** Other Candida strains used in this study.

| Species | Origin name | Number | Description |
| --- | --- | --- | --- |
| *C. albicans* | ATCC90028 | Clin01 | No known resistance |
| *C. tropicalis* | ATCC7349 | Clin03 | " " " |
| *C. parapsilosis* | ATCC22019 | Clin04 | " " " |
| *C. krusei* | ATCC6258 | Clin05 | " " " |
| *C. auris* | CBS10913 | Clin06 | " " " |
| *C. albicans* | CagS | Clin07 | " " " |
| *C. albicans* | CagR | Clin08 | Echinocandin resistant; serial isolate with Clin07 |
| *C. albicans* | DSY292 | Clin11 | Azole resistant |
| *C. albicans* | DSY296 | Clin12 | " " |
| *C. albicans* | DSY2323 | Clin15 | " " |
| *C. albicans* | CAAL74 | Clin16 | " " |
| *C. albicans* | AF-GIR | Clin17 | Echinocandin resistant |
| *C. albicans* | AF-ALA | Clin18 | " " |
| *C. albicans* | AF-DUB | Clin19 | " " |

**Figure S1.**
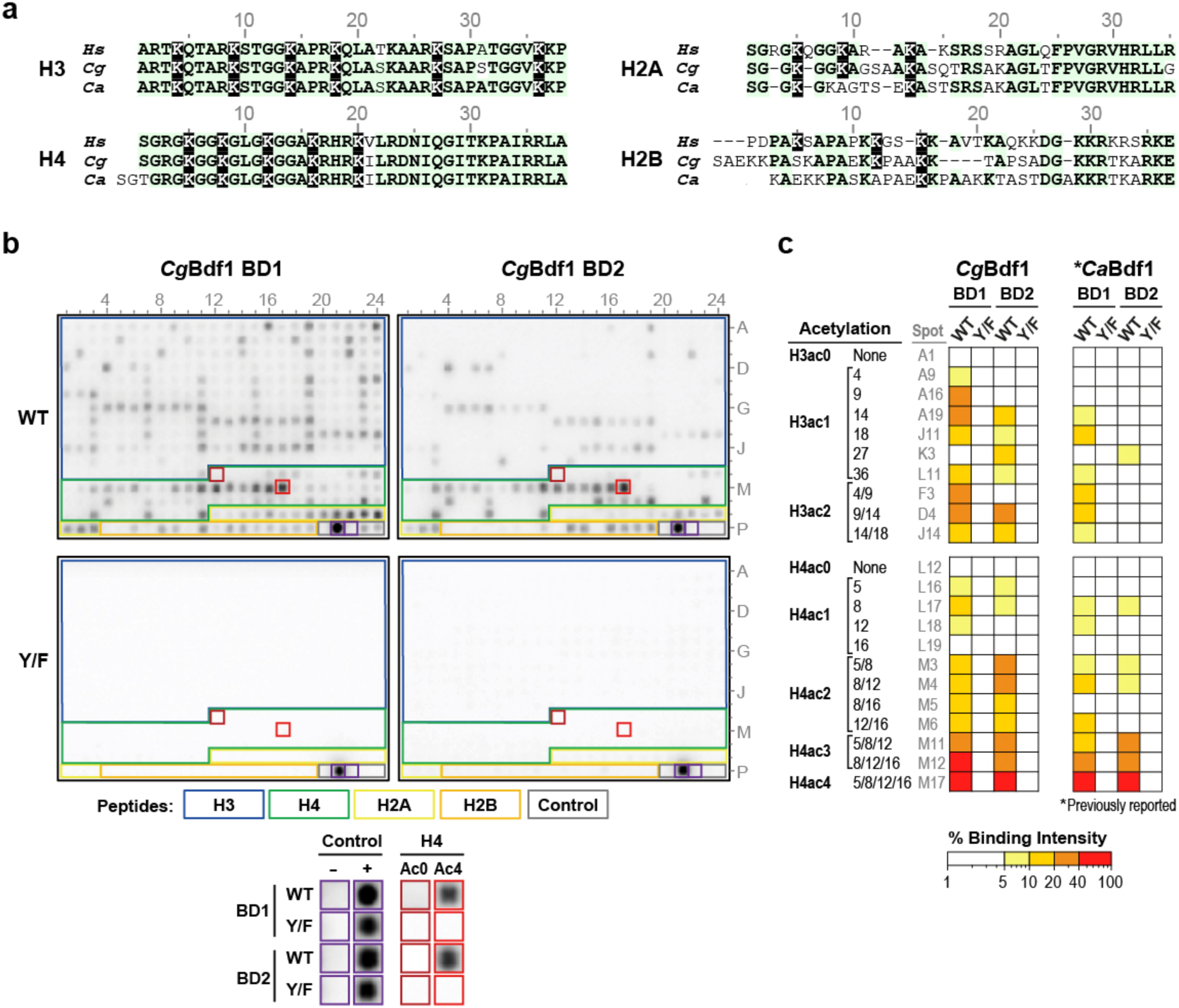
*Cg*Bdf1 BDs recognize multi-acetylated histone peptides. (**a**) Alignment of N-terminal sequences from human (*Hs*), *C. glabrata* (*Cg*) and *C. albicans* (*Ca*) histones showing the high sequence conservation of the H3 and H4 tails. Lysine positions acetylated in peptides on the array are shown in inverse font. (**b**) Binding of WT and mutant (Y/F) *Cg*Bdf1 BDs to a histone peptide array. The array includes control peptides (grey outline) and N-terminal peptides from histones H3, H4, H2A and H2B (blue, green, yellow and orange outlines, respectively), with H3 and H4 comprising the majority (347 of 384 peptides). Signals for background and positive-binding controls (boxed in red) or for unacetylated (Ac0) and tetra-acetylated (Ac4) H4 peptides (boxed in purple) are shown magnified below the array. (**c**) Summary of binding intensities for H3 and H4 peptides. The data for *Ca*Bdf1 BDs is from ref^29^.

**Figure S2.**
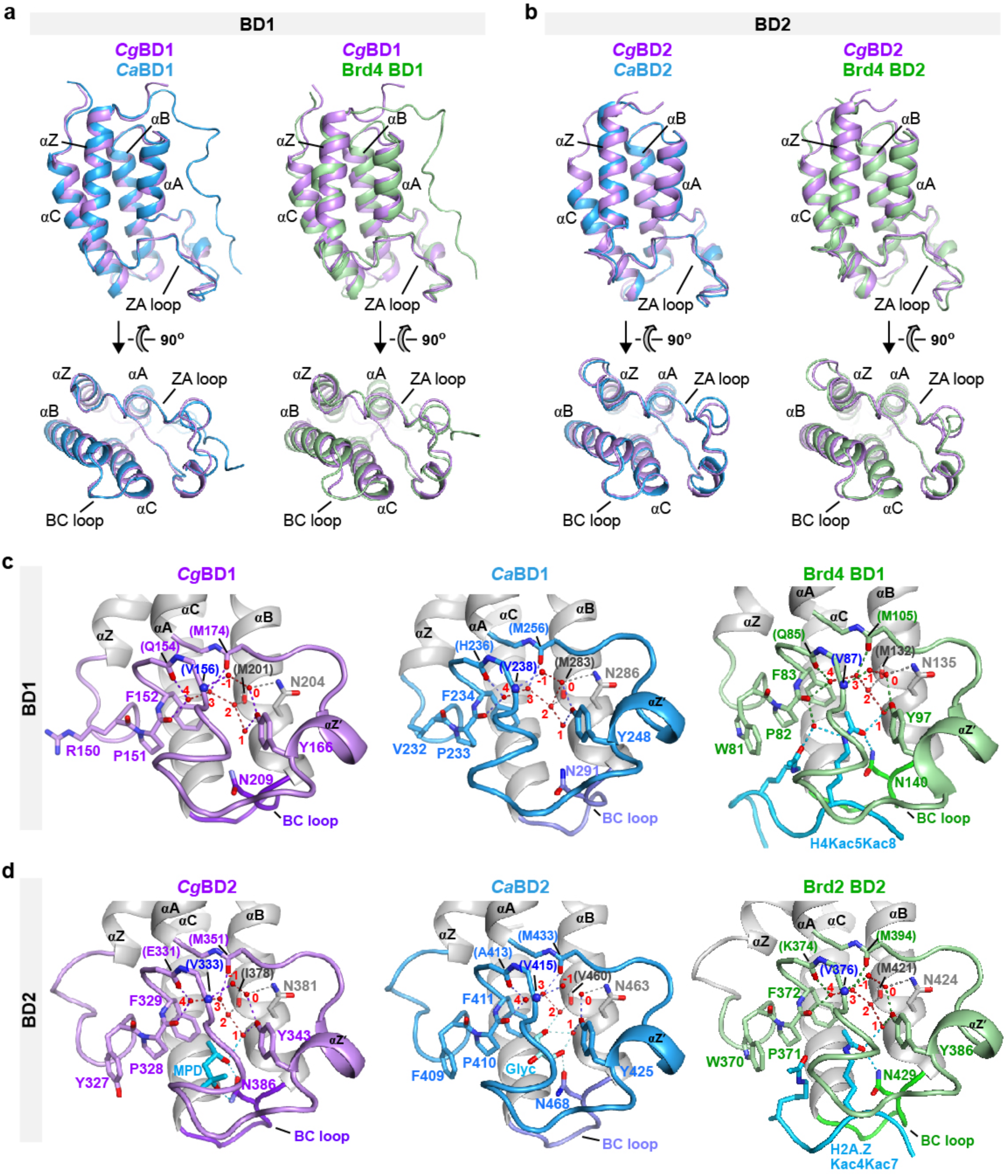
Structure of *Cg*Bdf1 BDs and comparison with *C. albicans* and human orthologs. **(a)** Alignment of the unbound BD1 structure from *Cg*Bdf1 (violet) with that of *Ca*Bdf1 (blue; PDB 5n15) and human Brd4 (green; PDB 2oss). **(b)** Alignment of the unbound BD2 structure from *Cg*Bdf1 (violet) with that of *Ca*Bdf1 (blue; PDB 5n18 chain B) and human Brd4 (green; PDB 2ouo). (**c**) Close-up view of *Cg*BD1 (violet) and *Ca*BD1 (blue, PDB 5n15) compared with that of human Brd4 BD1 bound to a diacetylated H4 peptide (green, PDB 3uvw) showing conserved water structure and hydrogen bonding interactions (dashed lines) in the ligand binding site. Residues interacting through backbone atoms are labelled in parentheses. Water molecules are numbered as in ref ^53^. (**d**) Ligand binding sites of *Cg*BD2 (violet) and *Ca*BD2 (blue) compared with that of human Brd2 BD2 bound to a diacetylated H2A.Z peptide (green, PDB 7jx7). Small molecules present in the crystallization buffers used for *Cg*BD2 (MPD) and *Ca*BD2 (glycerol) occupy the Kac binding pocket.

**Figure S3.**
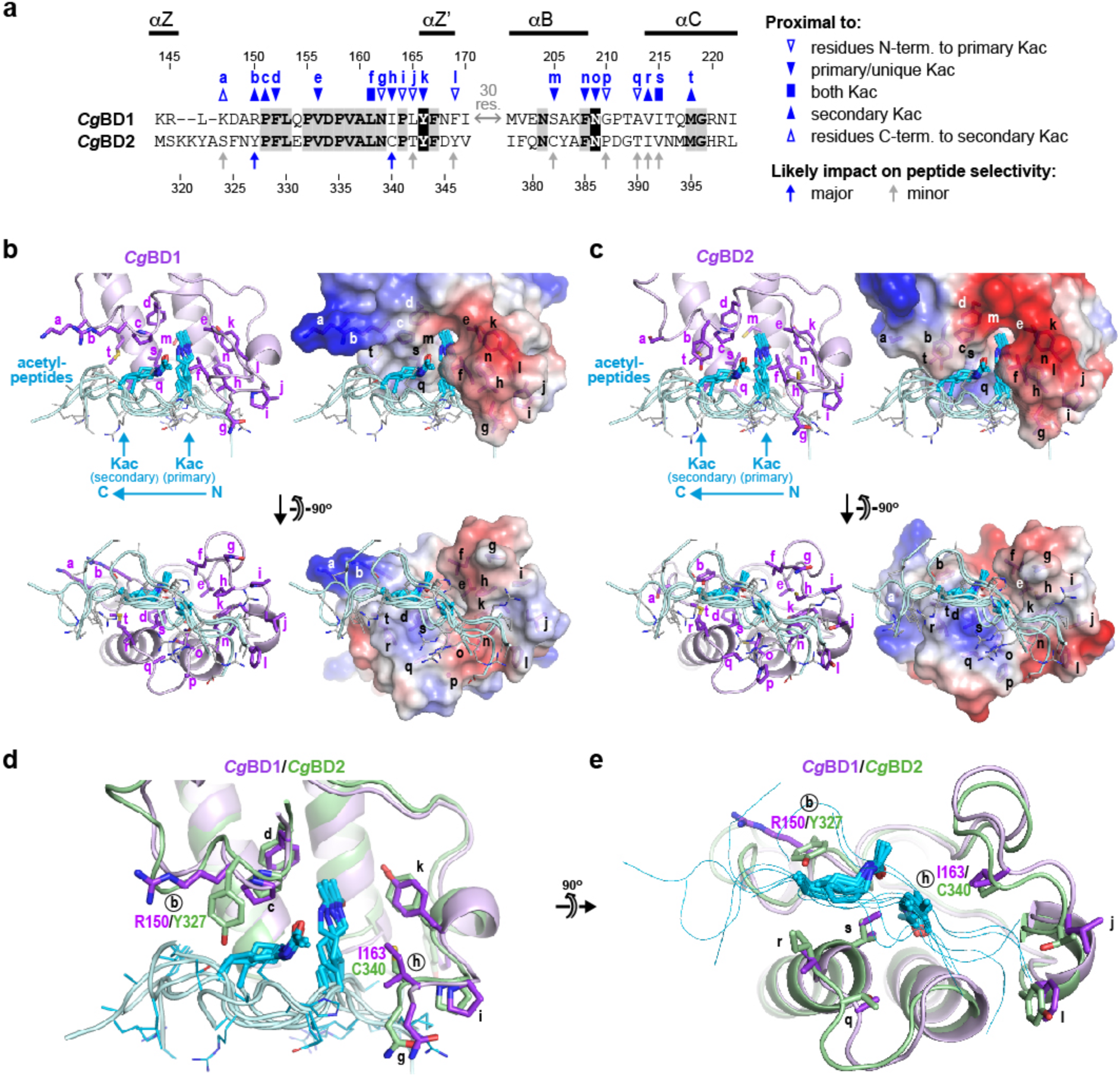
Comparison of the putative peptide-binding surfaces of *Cg*Bdf1 BD1 and BD2. (**a**) Sequence alignment of *Cg*BD1 and *Cg*BD2 residues in the ZA and BC loops and flanking regions. Residues labelled *a-t* correspond to human BET BD residues that line the peptide binding groove (defined as lying within 5 Å of an acetylated histone peptide) observed in the peptide-bound BD structures listed in (b). The 13 residue positions in these human BD structures that interact with one or both Kac residues are indicated by blue closed symbols, while positions contacting peripheral peptide residues are indicated by blue open symbols, as follows: down arrowheads indicate proximity to the primary/unique Kac residue (closed symbols) or to peptide residues N-terminal to this Kac (open symbols); up arrowheads indicate proximity to the secondary Kac (closed symbols) or to peptide residues C-terminal to this Kac (open symbols); a box indicates proximity to both Kac residues. Arrows beneath the alignment indicate the subset of residues *a*-*t* that differ between BD1 and BD2 and are hypothesized to have either a major (blue arrows) or at most only a minor (gray arrows) impact on peptide selectivity. The underlying reasoning is that residues poorly conserved between *Cg*BD1 and *Cg*BD2 and predicted to interact with the Kac residues are more likely to differentiate the ligand selectivity of these two BDs than are well-conserved residues that interact with the N- or C-terminal extremity of the bound peptide. (**b**) Structural alignment of *Cg*BD1 with peptide-bound human BET BD complexes. Unbound *Cg*BD1 was structurally aligned with the following human BET BDs bound to mono- or diacetylated histone tail peptides: Brdt BD1/H4K5acK8ac (PDB entry 2WP2), Brdt BD2/H3K18ac (2WP1), Brd4 BD1/H4K5acK8ac (3UVW), Brd4 BD1/H4K12acK16ac (3UVX), Brd4 BD1/H4K16acK20ac (3UVY), and Brd2/H2A.ZK4acK7ac (7JX7). (Entries 2WP2 and 2WP1 each had two BD/peptide complexes in the asymmetric unit, which were independently aligned). All peptides share a common binding mode in which the primary acetyllysine (AcK) residues bind deep within a central hydrophobic pocket, peptide residues N-terminal to the AcK interact with the B helix and C-terminal half of the ZA loop and peptide residues C-terminal to the AcK (including the secondary AcK) interact mainly with the C helix and N-terminal half of the ZA loop. For clarity, only the histone peptide component of the human BET complexes are shown. *Cg*BD1 is shown in light magenta and residues in the putative peptide-binding groove are shown as sticks and labeled *a-t* as in (a). Corresponding electrostatic surface representations are shown at right. Histone peptides are shown in cyan. The side chains for acK and non-acK residues are shown as thick and thin sticks, respectively. (**c**) Structural alignment of *Cg*BD2 with peptide-bound human BET complexes. (**d**) Structural alignment of *Cg*BD1 and *Cg*BD2 with histone peptides. Divergent residue pairs at positions *b* and *h* predicted to contribute to the minor differences in peptide selectivity observed between *Cg*BD1 and *Cg*BD2 are indicated by circled labels. (**e**) Orthogonal view to (d). For clarity, histone peptide backbones are represented as a thin trace and non-acK side chains are omitted.

**Figure S4.**
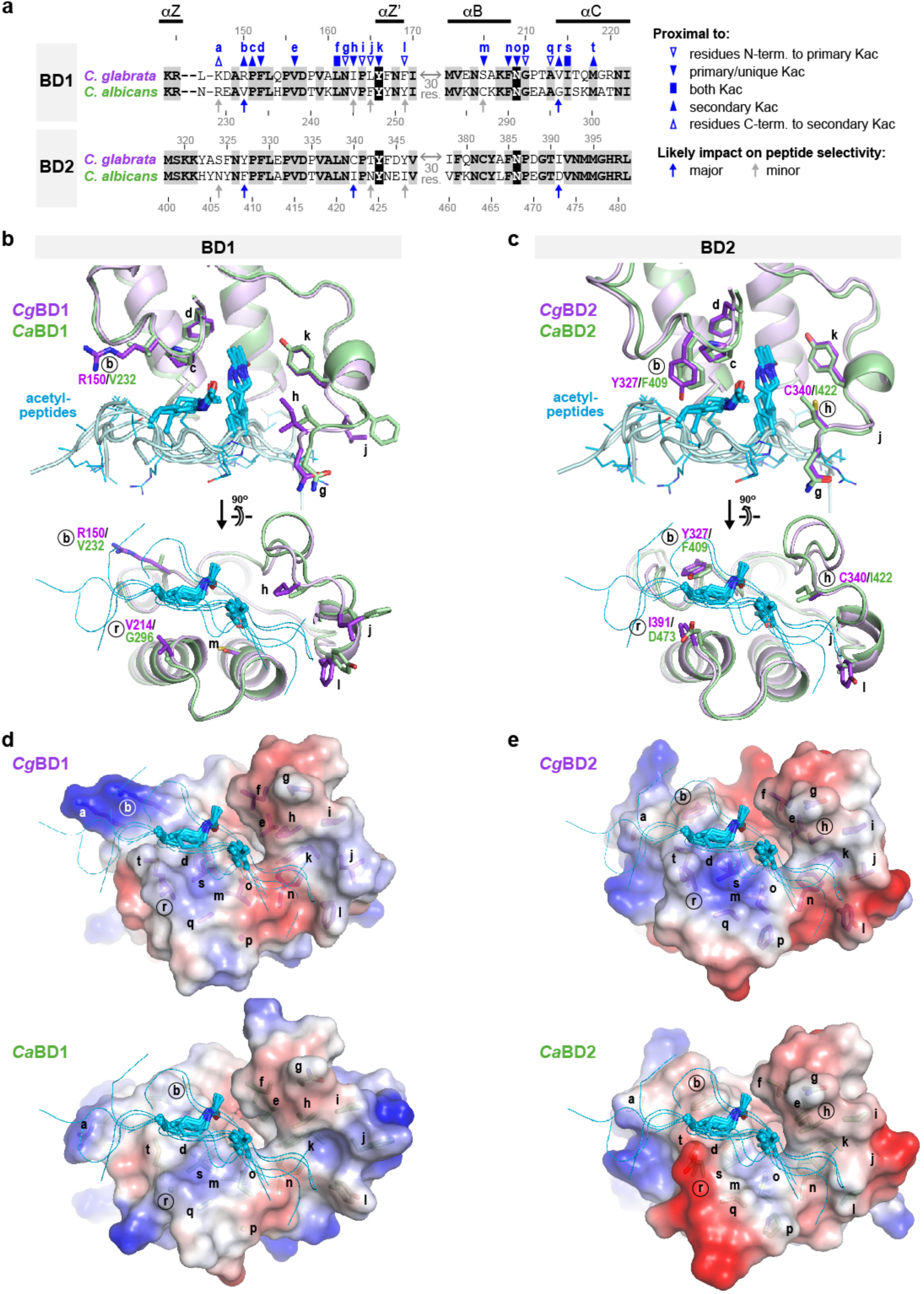
Comparison of the putative peptide-binding surfaces of *Cg*Bdf1 and *Ca*Bdf1 BDs. (**a**) Sequence alignment of the ZA and BC loops and flanking regions. Residue labels and symbols are as described in **Figure S3a**. (**b,c**) Structural alignment of (b) BD1 and (c) BD2 from *Cg*Bdf1 (violet) and C*a*Bdf1 (green) with peptide-bound human BET BD complexes. (Human BDs used in the alignment are listed in **Figure S3b**.) For clarity, only the histone peptide component (cyan) of the human BD complexes is shown. Peptide side chains for acK residues are shown as thick sticks while those for non-acK residues are shown as thin sticks (upper panels) or are omitted (lower panels). A subset of BD residues in the putative peptide-binding groove are shown as sticks and labeled with lower-case letters as in (a). Circled labels indicate residue positions predicted to contribute significantly to differential peptide selectivity. (**d,e**) Electrostatic surface representations of (d) BD1 and (e) BD2 from *Cg*Bdf1 (top) and *Ca*Bdf1 (bottom). In *Ca*BD2, residue D473 at position *r* confers a negative charge in the vicinity of the Kac binding site that is absent from the other three BDs. Together with substitutions at positions *b* and *h* this may explain why *Ca*BD2 binds a narrower range of acetylated H3 and H4 peptides compared to those of *Cg*BD1*, Cg*BD2 and *Ca*BD1.

**Figure S5.**
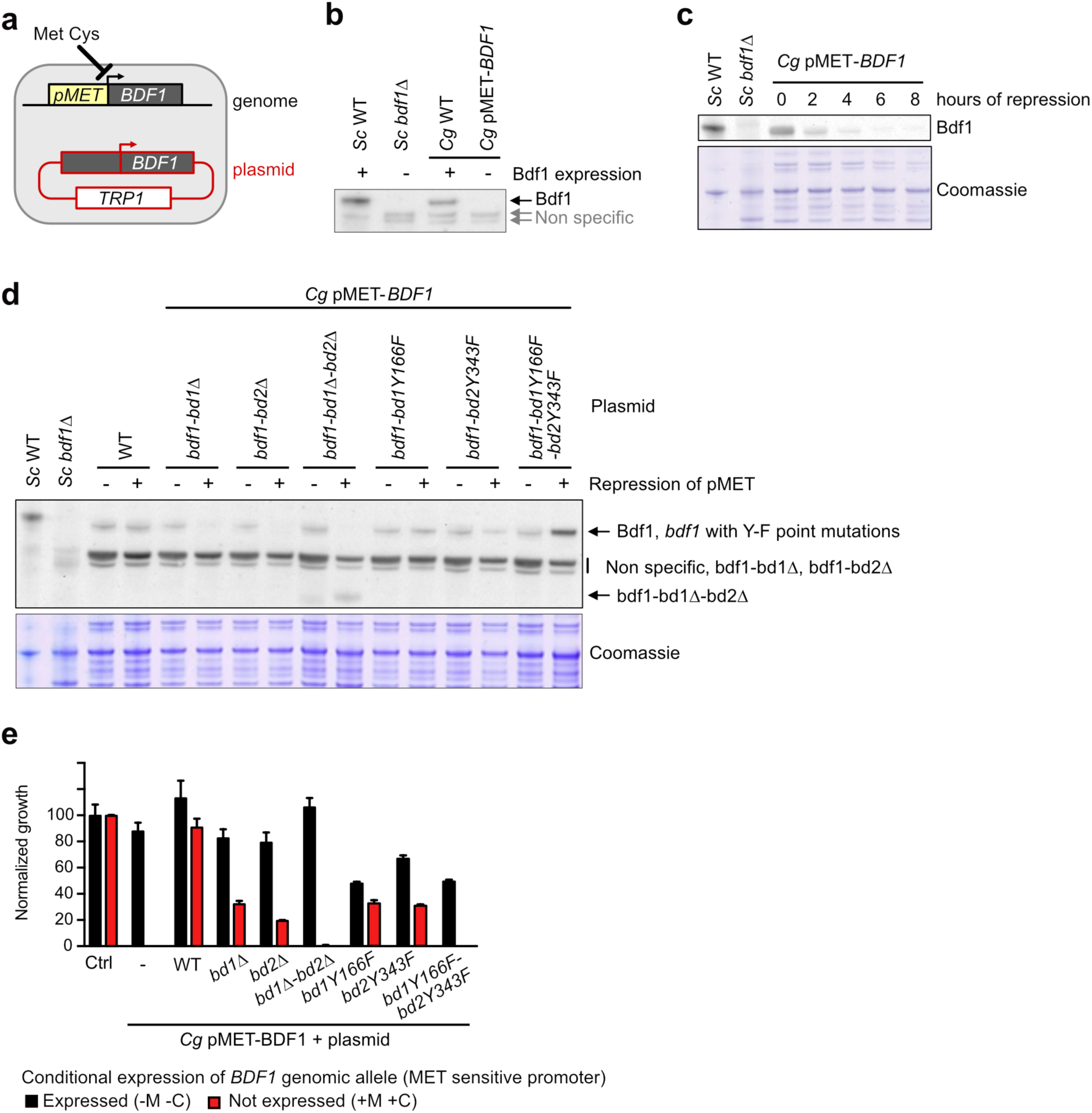
*Cg*Bdf1 bromodomains are essential for yeast viability. (**a**) Schematic of the genetic system. The endogenous *BDF1* promoter was replaced by a *MET3* promoter (pMET), which is repressed in the presence of cysteine and methionine. *BDF1* expression is rescued from a autonomous plasmid. (**b**) Validation of the Bdf1 antibody ^27^, which cross-reacts between Bdf1 from *S. cerevisiae* (Sc) and *C. glabrata* (Cg). (**c**) Repression of *BDF1* expression following the addition of methionine and cysteine. (**d**) Expression levels of *BDF1* upon repression of its genomic copy. Its expression is rescued from a plasmid (*BDF1* WT or mutant). (**e**) Impact of repression of the genomic copy of *BDF1* on vegetative growth with ectopic expression of WT or mutant *BDF1* from a plasmid.

**Figure S6.**
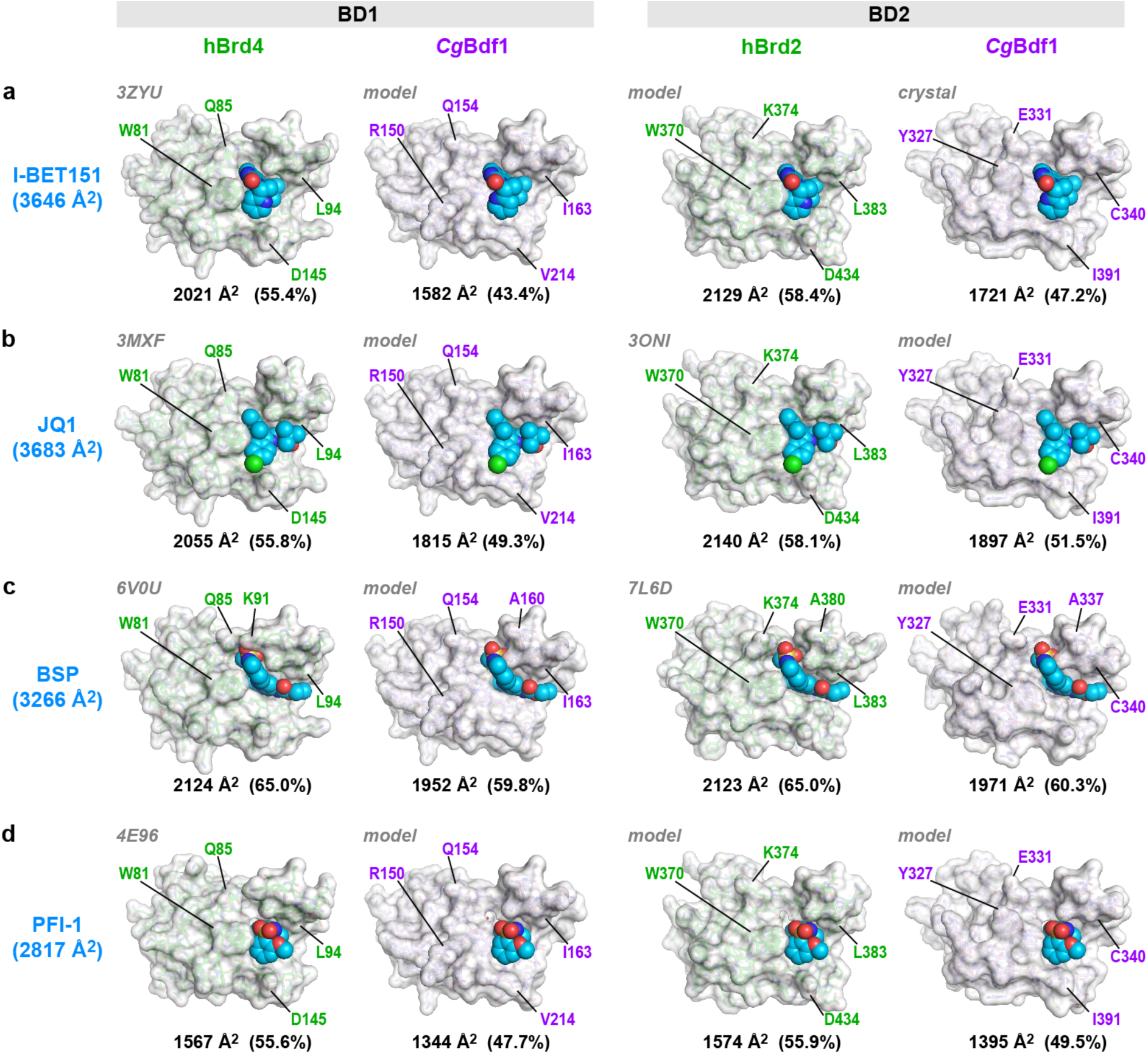
A wider binding pocket explains the lower potency of BET inhibitors toward *Cg*Bdf1 BDs. Crystal structures and hypothetical models of human BET and *Cg*Bdf1 BDs bound to (**a**) I-BET151, (**b**) JQ1, (**c**) bromosporine (BSP) and (**d**) PFI-1. The total solvent-accessible surface area (SASA) of the isolated BETi compound is shown in cyan. The compound’s SASA that is buried by each BD is indicated below the structure. Crystal structures are shown for human Brd4 BD1 bound to I-BET151, JQ1, BSP and PFI-1, human Brd2 BD2 bound to JQ1 and BSP (PDB entries 3ZYU, 3MXF, 6V0U, 4E96, 3ONI and 7L6D, respectively) and *Cg*BD2 bound to I-BET151 (this work). Brd2 BD2 bound to I-BET151 and PFI-1 were modelled by aligning the corresponding ligand-bound structure of Brd4 BD1 onto that of JQ1-bound Brd2 BD2 and replacing the atomic coordinates of JQ1 with those of I-BET151 or PFI-1. BETi-bound *Cg*BD1 structures were modelled by aligning the structure of *Cg*BD2 bound to I-BET151 or that of Brd4 BD1 bound to JQ1, BSP or PFI-1 onto the unbound *Cg*BD1 crystal structure, altering the *Cg*BD1 residue Ile215 rotamer to match the corresponding Brd4 Ile146 rotamer (to avoid a steric clash with the ligand), and transferring the ligand coordinates. Structure alignments predict that JQ1 and PFI-1 are sterically incompatible with the *gauche*^-^ conformation of *Cg*BD2 Tyr327 but that BSP is compatible with both the *trans* and *gauche*^-^ conformations. Accordingly, JQ1- and PFI-1-bound *Cg*BD2 were modelled by aligning the corresponding Brd4 BD1 structure onto that of I-BET151-bound *Cg*BD2 and replacing the I-BET151 coordinates with those of JQ1 or PFI-1, whereas BSP-bound *Cg*BD2 was modelled by aligning the BSP-bound structure of Brd4 BD1 onto that of unbound *Cg*BD2 and transferring the BSP coordinates. All ions, non-BETi ligands and water molecules (except for the 6 structurally conserved waters, numbered -1 to 4 in **Figure SBc** and **d**) were removed from each structure prior to calculating SASA values in PyMOL ^56^.

**Figure S7.**
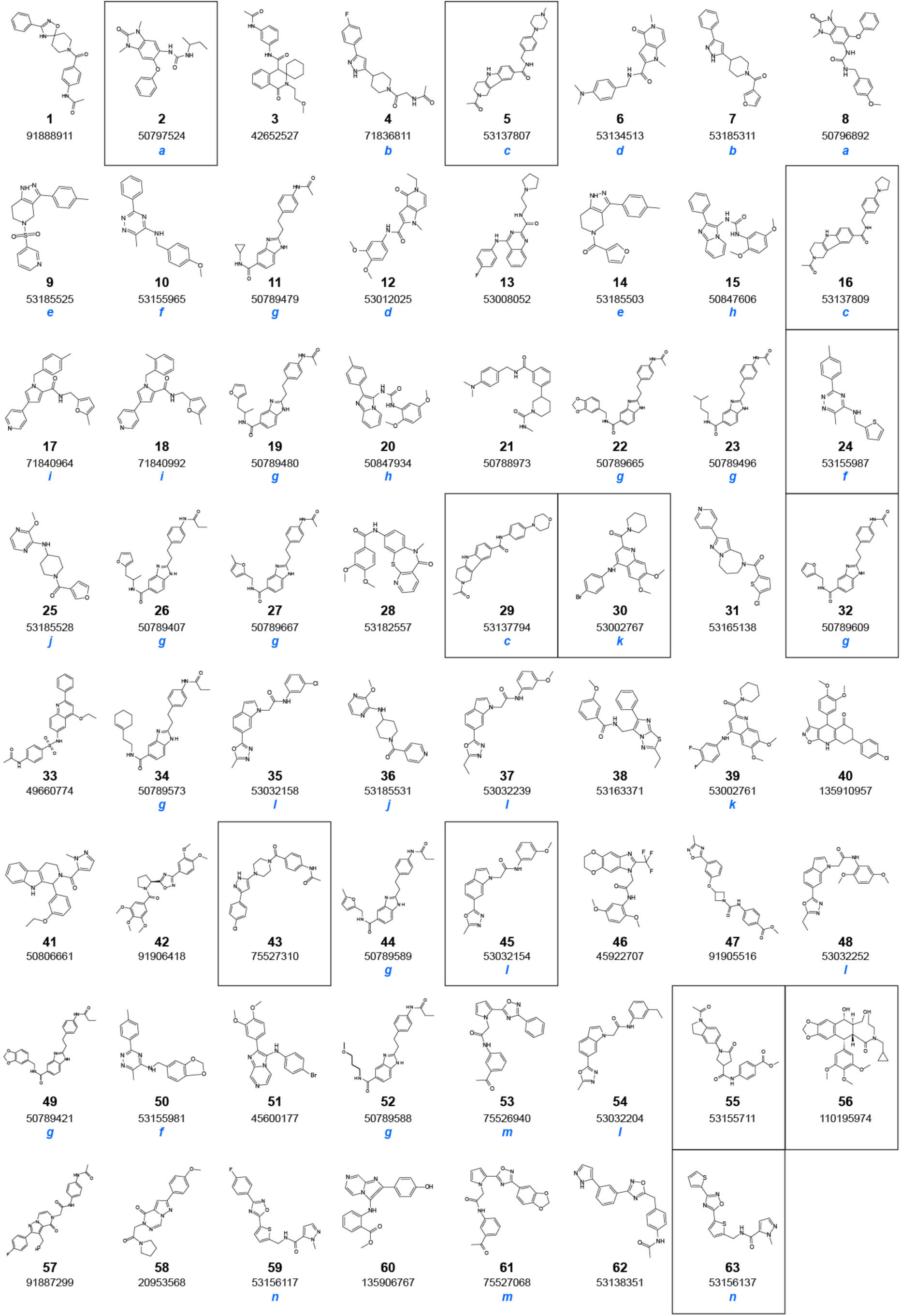
*Cg*Bdf1 BD1 inhibitors identified by high-throughput screening. The PubChem Compound ID is indicated for each inhibitor. Compounds sharing the same scaffold are labeled with the same letter (*a-n*) in blue. The 12 compounds pursued for detailed characterization are boxed.

**Figure S8.**
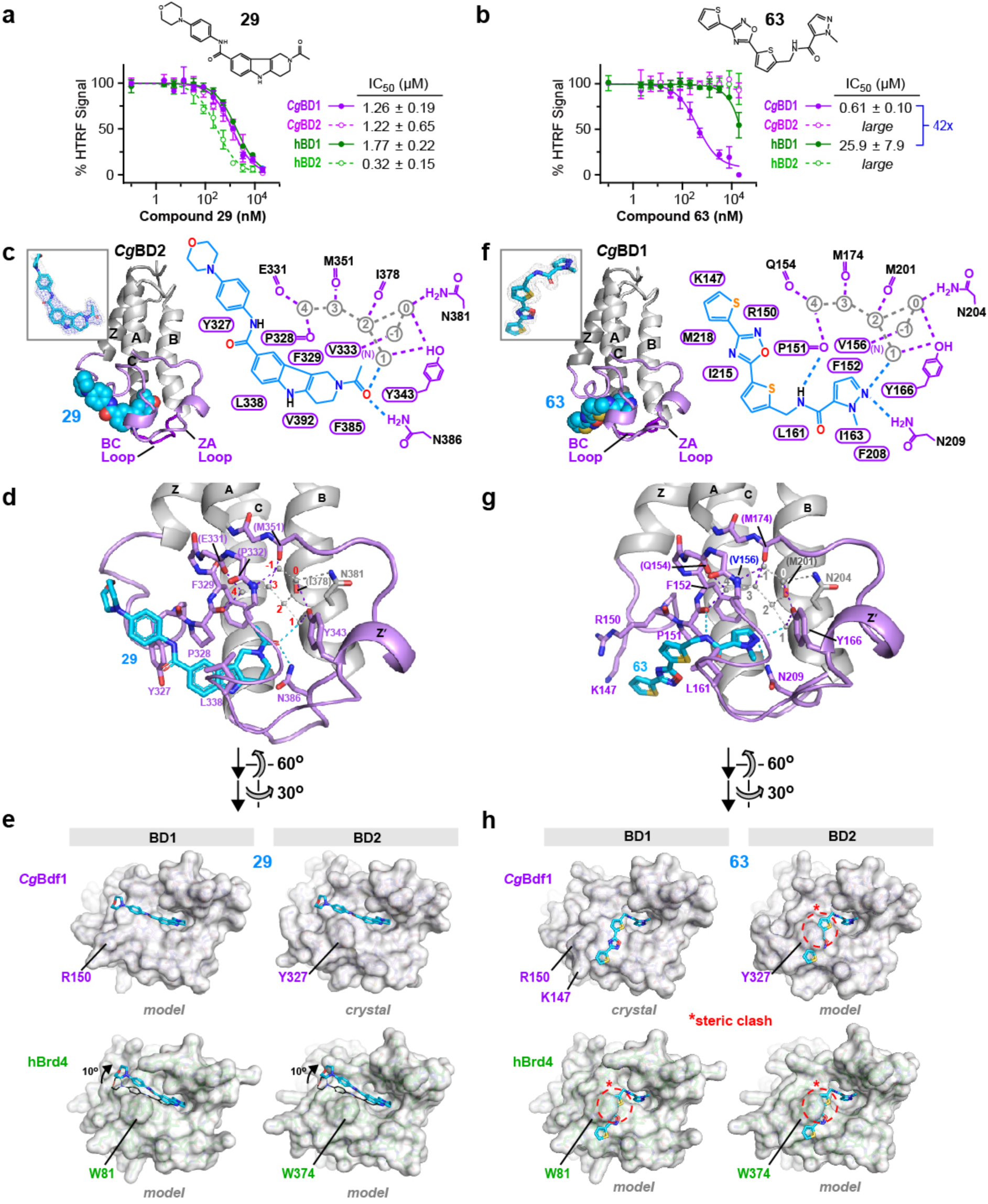
Structural basis of non-selective and selective inhibition of *Cg*Bdf1 BDs by 29 and 63. (**a**,**b**) HTRF assays performed on *Cg*Bdf1 BD1 and BD2 (closed purple and open magenta circles, respectively) and human Brd4 BD1 and BD2 (closed dark green and open light green circles, respectively) in the presence of compound (a) **29** and (b) **63**. (**c**) Crystal structure of *Cg*BD2 bound to **29** and schematic summary of ligand interactions. Hydrogen bonds are shown as dashed lines. Residues mediating van der Waals contacts with I-BET151 are indicated by labels within a cartouche. Water molecules are indicated in grey and are numbered as in ref. ^53^. *Inset.* Simulated annealing omit *F*_o_-*F*_c_ density for **29** contoured at 3 *σ*. **d**) Details of the ligand binding site. Residues interacting with **29** through direct and water-mediated hydrogen bonds (dashed lines) are shown in stick representation. Residues interacting through backbone atoms are labelled in parentheses. (**e**) Surface representations of the *Cg*BD2/**29** crystal structure and structural alignment models of *Cg*BD1, Brd4 BD1 and BD2 showing how the four BD binding pockets could accommodate **29**. The presence of WPF Trp residues 81 and 374 in Brd4 is predicted to induce a small rotation of **29** relative to its orientation in CgBD2. (**f**) Crystal structure of *Cg*BD2 bound to **29** and schematic summary of ligand interactions. *Inset.* Simulated annealing omit *F*_o_-*F*_c_ density for **63** contoured at 3 *σ*. (**g**) Details of the ligand binding site. Residues interacting with **63** through direct and water-mediated hydrogen bonds (dashed lines) are shown in stick representation. Residues interacting through backbone atoms are labelled in parentheses. (**h**) Surface representations of the *Cg*BD1/**63** crystal structure and structural alignment models of *Cg*BD2, Brd4 BD1 and BD2 showing observed or predicted interactions with the binding pocket. The steric clashes predicted between **63** and residues Tyr327 of *Cg*BD1, Trp81 of Brd4 BD1 and Trp374 of Brd4 BD2 are indicated by red circles and asterisks.

